# Validating the potential of double-stranded RNA targeting Colorado potato beetle *mesh* gene in laboratory and field trials

**DOI:** 10.1101/2020.02.13.945097

**Authors:** Marko Petek, Anna Coll, Rok Ferenc, Jaka Razinger, Kristina Gruden

**Affiliations:** National Institute of Biology, Department of Biotechnology and Systems Biology, Ljubljana, Slovenia; Agricultural Institute of Slovenia, Plant Protection Department, Ljubljana, Slovenia

**Keywords:** RNA interference (RNAi) feeding, dsRNA, gene silencing, RNAi pest control, survival analysis, *Leptinotarsa decemlineata* (Say), field trial, *E coli* HT115 (DE3)

## Abstract

Colorado potato beetle (CPB) is an agricultural pest of solanaceous crops, notorious for its rapid resistance development to chemical pesticides. Foliar spraying of dsRNA formulations is a promising innovative technology providing highly specific and environmentally acceptable option for CPB management.

We designed dsRNA to silence CPB *mesh* gene (dsMESH) and performed laboratory feeding trials to assess impacts on beetle survival and development. We compared the effectiveness of *in-vivo* and *in-vitro* produced dsRNA in a series of laboratory experiments. We additionally performed a field trial in which the efficacy of dsRNA sprayed onto potato foliage was compared to a spinosad-based insecticide.

We showed that dsMESH ingestion consistently and significantly impaired larval growth and decreased larval survival in laboratory feeding experiments. *In-vivo* produced dsRNA performed similarly as *in-vitro* synthesised dsRNA in laboratory settings. In the field trial, dsMESH was as effective in controlling CPB larvae as a commercial spinosad insecticide, its activity was however slower. We discuss limitations and benefits of a potential dsMESH-based CPB management strategy and list some important RNAi based CPB research topics, which will have to be addressed in future.

## INTRODUCTION

Colorado potato beetle (CPB), *Leptinotarsa decemlineata*, is a serious pest of potato and other solanaceous crops. It is well known for its ability to rapidly evolve resistance to insecticides; it has already evolved resistance to all major insecticide classes (Alyokhin et al. 2008). Extensive use of conventional insecticides can have undesirable effects on the environment, non-target organisms and human health. Compared to chemical pesticides, double-stranded RNAs (dsRNAs) have the advantage of high selectivity towards the target organism and rapid environmental degradation into non-toxic compounds (Dubelman et al. 2014; Albright et al. 2017). Therefore, this novel pest management approach has the potential to decrease the extensive use of conventional insecticides.

When delivered into cells, dsRNAs activate the RNA interference (RNAi) mechanism that mediates a sequence-specific suppression of transcription, also called gene silencing (Joga et al. 2016). In CPB, unlike some other insects, dsRNAs are not degraded by gut nucleases, are efficiently taken up by the gut epithelium cells, and can trigger local as well as systemic RNAi response (Cappelle et al. 2016). This makes CPB an excellent candidate for pest management using dsRNAs, which was first recognized in a study by Baum et al. (2007) that has also identified several RNAi targets. This was followed by studies that identified novel effective target genes in CPB (Zhu et al. 2011, 2015; Zhou et al. 2013; Wan et al. 2014; Meng et al. 2015, 2018; Lü et al. 2015; Fu et al. 2015, 2016; Shi et al. 2016a, b; Guo et al. 2016; Xu et al. 2018) and for some targets also validated in field trials (Guo et al. 2018). In western corn rootworm, Hu et al. (2016) identified another target gene – *mesh* (alternatively named *dvssj2*) which encodes a smooth septate junction protein important for structural integrity of the midgut epithelium. They showed that silencing *mesh* impairs midgut barrier function which results in increased larval mortality (Hu et al. 2019).

In this study, we used *in vitro* and *in vivo* synthesised dsRNA designed to silence the *mesh* gene in CPB. We performed laboratory-based feeding assays with CPB at different stages of larval development as well as a small-scale field trial to validate the designed dsRNA’s pesticidal potential in a commercial production system. Therefore, our study offers new data on dsMESH effectiveness in another coleopteran, CPB, which represents an important crop pest and the most probable first candidate for spray-induced gene silencing commercialization.

## MATERIALS AND METHODS

### Quantification of *mesh* gene expression by qPCR

To quantify the expression levels of *mesh*, RNA was extracted from three to four individual larvae (three to four biological replicates), except for the study of *mesh* expression in CPB body parts where one pooled sample from 3-4 beetles for each body part was analysed. RNA extraction was performed using TRIzol reagent (Invitrogen) and Direct-zol RNA Microprep kit (Zymo Research). DNase treatment and reverse transcription were performed as described previously (Petek et al. 2012). RNA concentration and integrity were validated using a NanoDrop ND-1000 spectrophotometer and agarose gel electrophoresis. The efficiency of DNase treatment was confirmed by qPCR with no RT samples.

The expression of *mesh* was assessed by quantitative PCR (qPCR). *Mesh* gene model from i5k genome version 0.5.3 (LDEC006484; Schoville et al. 2018) was corrected based on alternative models and mapped RNA-seq reads available in i5k’s WebApollo instance (Supplementary Figure 1 and Supplementary Data 1.1). The qPCR primers and probes were designed in Primer Express 2.3 (Applied Biosystems) using default parameters for TaqMan amplicons and were synthesised by IDT. Assay specificity was verified *in silico* using blastn queries against all transcripts predicted in the CPB genome (Schoville et al. 2018). The linear ranges and amplification efficiencies were determined across five 10-fold serial dilutions of cDNA. Target gene accumulation was normalised to three endogenous control genes: LdRP4 (Shi et al. 2013), 18S rRNA (Eukaryotic 18S rRNA Endogenous Control, Applied Biosystems) and LdSmt3 (Petek et al. 2014). Primer and probe sequences, qPCR chemistry and other assay metadata are available in Supplementary Table 1.

FastStart Universal Probe Master Rox mastermix (Roche) was used for TaqMan chemistry based assays and Power SYBR mastermix (Applied Biosystems) for SYBR Green chemistry based assays. Dilution of cDNA samples and pipetting of qPCR reagents onto 386-well plates was performed on a Microlab STARlet automated liquid handling system (Hamilton). Reactions were performed in 5 μl total volume on LightCycler 480 (Roche) as described previously (Petek et al. 2012). Melting curve analysis was applied for SYBR green chemistry based assays LdRP4 and LdSmt3 to control for primer dimer formation and amplification specificity in each reaction. Each sample was analysed in two replicates of two dilutions to check for the presence of inhibitors in the sample. Cq values were calculated using instrument manufacturer software and exported as text files. Amplification quality control for each sample and relative quantification based on the standard curve method was performed in quantGenius software (Baebler et al. 2017). For every gene, the limit of quantification (LOQ) was determined from the standard curve. The normalized target copy numbers calculated by quantGenius were exported to an Excel file to calculate standard errors and Student’s *t*-test statistics.

### Design and *in vitro* synthesis of dsRNAs

To avoid sequence regions that might affect other species due to nucleotide conservation we used EMBOSS splitter (Rice et al. 2000) to generate all possible 21-mers for the CPB *mesh* transcript. These 21-nt sequences were queried using BLASTn against non-target organism transcriptomes including *Homo sapiens, Apis mellifera, Bombus terrestris, Danaus plexippus, Drosophila melanogaster, Megachile rotundata, Nasonia vitripennis*, and *Tribolium castaneum*. Regions of the transcripts with 20 or 21 nt BLASTn hits in non-target organisms were excluded from dsRNA design. Based on the above metrics, the longest CPB-specific region was selected as the input sequence to design a long dsRNA molecule using e-RNAi web service (Horn and Boutros 2010) using default parameters. Such bioinformatics design however does not exclude the possibility of off-target effects. For example, due to crosstalk between siRNA and miRNA pathways, off-target silencing could be triggered by siRNAs with less sequence conservation. Also, due to limited genomics and transcriptomics sequence availability in Arthropods a comprehensive bioinformatics analysis is not possible (Christiaens et al. 2018). As non-specific dsRNA control, the dsEGFP with sequence corresponding to a fragment of enhanced green fluorescent protein (Guo et al. 2015) was used (sequences in Supplementary Data 1.2). *In vitro* synthesis of dsMESH and dsEGFP was performed by AgroRNA (South Korea). The quality and quantity of dsRNA was determined using agarose gel electrophoresis and NanoDrop.

### *In vivo* synthesis of dsRNAs

To *in vivo* synthesise dsMESH, a 417 bp fragment of the gene was amplified by PCR from a pooled CPB midgut cDNA sample using Phusion DNA polymerase (Biolabs) and cloned into L4440gtwy (Addgene) using pCR8/GW/TOPO TA Cloning Kit (Invitrogen) to obtain MESH::L4440. The correct fragment insertion was confirmed by Sanger sequencing (Eurofins Genomics). The GFP::L4440 plasmid (Addgene), containing a full-length (857 bp) green fluorescence protein sequence insert was used to produce dsGFP. Heat-shock induced competent *Escherichia coli* HT115 (DE3) bacteria were transformed with MESH::L4440 and GFP::L4440, respectively. Transformation was confirmed by colony PCR using KAPA2G Robust HotStart Polymerase (Kapa Biosystems).

To produce dsRNA, cultures of transformed bacteria were grown to OD600 0.5 in 250 ml of liquid LB media. Production of dsRNAs was induced with 400 μM IPTG (Thermo Scientific). After 4 h, cells were pelleted, re-suspended in 6 ml nuclease-free water (Sigma) and lysed by boiling followed by four freezing-thawing cycles and a 15 min treatment in ultrasonic bath SONIS 4 (Iskra PIO). Bacterial lysates were centrifuged at 9000 g for 20 min and the supernatant was concentrated to 1/10 volume using GeneVac EZ-2plus (Genevac Ltd).

To estimate the quantity of dsRNA produced, total RNA was extracted from bacterial lysates using Direct-zol RNA MiniPrep Plus kit (Zymo Research), treated with DNase I (Zymo Research) and reverse transcribed using High-Capacity cDNA Reverse Transcription Kit (Applied Biosystems). *In vivo* synthesised dsMESH and dsGFP quantities were estimated from 1% agarose E-Gel EX (Thermo) RNA band intensities. Identity of dsRNA was confirmed by RNase If treatment (Supplementary Data 1.3, Supplementary Figure 2).

### Laboratory feeding trials

CPBs were reared on potato plants cv. Désirée in conditions described previously (Petek et al. 2014). Larvae, which hatched on the same day, were reared on non-treated detached potato leaves until most larvae reached desirable treatment stage. For each feeding trial, larvae were randomly selected and assigned into treatment groups, enclosed into plastic or glass containers and reared on untreated potato foliage. DsRNA were either sprayed on detached leaves, potted whole plants, or CPB eggs, or pipetted onto freshly cut leaf disks (Table 1). To protect detached leaves from desiccation, the petioles were placed in sterile 2 ml microcentrifuge tubes filled with 0.5% agarose gel, whereas leaf disks were placed into flat bottom 24-well plates with bottom covered by 0.5% agarose gel. After consumption of leaf disk (trials three and four, Table 1), the larvae were moved back to plastic containers and daily supplied with untreated detached potato leaves.

**Table 1.** Design of Colorado potato beetle dsRNA laboratory-based feeding trials.

| Feeding trial number | Conducted | Negative controls | dsRNA production | CPBs per treatment | CPB stage at first treatment | Treatment regime (dsRNA spray concentration [serial dilution]; dose per larva) | Trial duration (d) |
| --- | --- | --- | --- | --- | --- | --- | --- |
| 1 | Jun-Jul 2016 | water, dsEGFP | *in vitro* | 40 | 2<sup>nd</sup> instar larvae | continuous feeding on sprayed detached leaves (conc. 0.4 µg/µl) | 41 |
| 2 | Jun-Jul 2016 | water, dsEGFP | *in vitro* | 30 | 4<sup>th</sup> instar larvae | continuous feeding on potted plants sprayed once (conc. 0.4 µg/µl) | 24 |
| 3 | Dec 2016 | water, dsEGFP | *in vitro* | 16 | 2<sup>nd</sup> instar larvae | discontinuous feeding on leaf disks (conc. 0.5 µg/µl; 0.75 µg/larva) | 7 |
| 4 | Apr 2018 | water, dsGFP (*in vivo*) | *in vitro*,<br>*in vivo* | 24 | 2<sup>nd</sup> instar larvae | discontinuous feeding on leaf disks (*in vitro*: conc. 0.1 µg/µl [60x], 0.01 µg/µl [600x], 0.001 µg/µl [6000x]; dose 0.6 µg/larva, 0.06 µg/larva, 0.006 µg/larva, respectively) (*in vivo*: conc. 0.1 µg/µl; dose 0.6 µg/larva) | 10 |
| 5 | May 2018 | water, dsEGFP | *in vitro* | 20 egg masses | eggs | egg spraying (conc. 0.5 µg/µl) | 7 |

Nuclease-free water (Sigma) was used for blank control treatment and dilution of all dsRNAs. In all, except trial four, *in vitro* synthesised dsEGFP (Guo et al. 2015) was used as non-specific dsRNA treatment control. In trial four, *in vivo* synthesised dsGFP sequence from GFP::L4440 plasmid (Addgene) was used instead. During the study, we adhered to national and institutional biosafety standards. More details on feeding trials are given in Table 1 and Supplementary Data 1.4-8.

Analysis of right-censored survival data was performed using Cox proportional hazards regression model fit and statistical tests implemented in R survival package version 2.42 (Therneau and Grambsch 2000). Data analysis execution calls are given in Supplementary Data 1.9.

### Field trial

A small-scale field trial was conducted in June and July 2019 on three locations near Ljubljana, Slovenia. Two trials were performed on adjacent potato fields in Šentjakob (46°05’13.4”N 14°34’06.8”E) and one in Iška vas (45°56’28.8”N 14°30’31.0”E). The experiment was designed following EPPO guidelines (EPPO 2008). Cultural conditions were uniform for all plots of the trial at each location and conformed to local agricultural practice. To assess the efficacy of dsMESH the only difference between treatments was the method of CPB management. Three 25 m^2^ plots were marked at each location. Each plot was divided into four replicate sub-plots, giving four replicates per treatment. On each sub-plot, an individual potato plant infested by at least 15 CPB larvae was randomly selected and marked, giving four plants per treatment at each location. Before treatment, foliage from potato plants and weeds surrounding the marked potato plants was removed to restrict larval movement between plants. Any unhatched CPB eggs from the marked potato plants were removed. CPB larvae were counted and larval stages and plant defoliation percentages were determined for each plant separately. The leaf damage caused by the CPB larval herbivory was estimated visually by inspecting the first ten fully developed leaves from the topmost apical plant meristem downwards on each marked potato plant.

Marked plants were sprayed with *in-vitro* synthesised dsMESH in concentration 10 μg/ml. We used potato plants sprayed with the manufacturer recommended 0.5% diluted spinosad formulation (insecticide Laser 240 SC, Dow AgroSciences) as a positive control treatment and unsprayed plants as a negative control. Two days post treatment (dpt), CPB larvae on marked plants were recounted and stages determined, and 7 dpt larvae were counted again, their stages determined, and leaf damage estimated. From the relative change of leaf damage assessed before treatment and at 7 dpt, the parameter ‘leaf damage increase’ was calculated.

Statistical differences in leaf damage, leaf damage increase, and larval mortality according to Henderson-Tilton were calculated using ANOVA and Bonferroni’s Multiple Comparison Test in GraphPad Prism 5.00 (GraphPad Software). The dataset was also analysed using a general linear model (GLM), where the effect of factors *treatment* (dsMESH, spinosad, and control), *experiment* (trials 1-3) and *replicate* (1-4) on previously mentioned parameters was assessed. Further, Fisher’s least significance difference (LSD) procedure at 95% confidence level was used to discriminate between the treatments within the three-trial dataset. These analyses were performed with the statistical software Statgraphics Centurion XVI (StatPoint Technologies).

## RESULTS

### The target gene *mesh* is expressed throughout all CPB developmental stages

To test whether an RNAi insecticide targeting *mesh* will work against all CPB’s developmental stages, we profiled *mesh* expression through the stages. Constitutive expression of *mesh* was detected in all developmental stages. Expression in the gut is highest in fourth instar larvae preceding pupation and in adults (Figure 1 and Supplementary Dataset 1). Constitutive expression of *mesh* in larval and adult stages is also evident from mapped RNA-Seq data available at i5k CPB genome browser (Supplementary Figure 1A). This expression pattern is suitable for RNAi insecticide targets as the gene is expressed in stages in which the beetles feed on plant leaves. We also qualitatively showed higher expression of *mesh* in samples of foregut, midgut and hindgut tissues compared to samples of legs, head and antennae (Supplementary Figure 3 and Supplementary Dataset 2).

**Figure 1.**
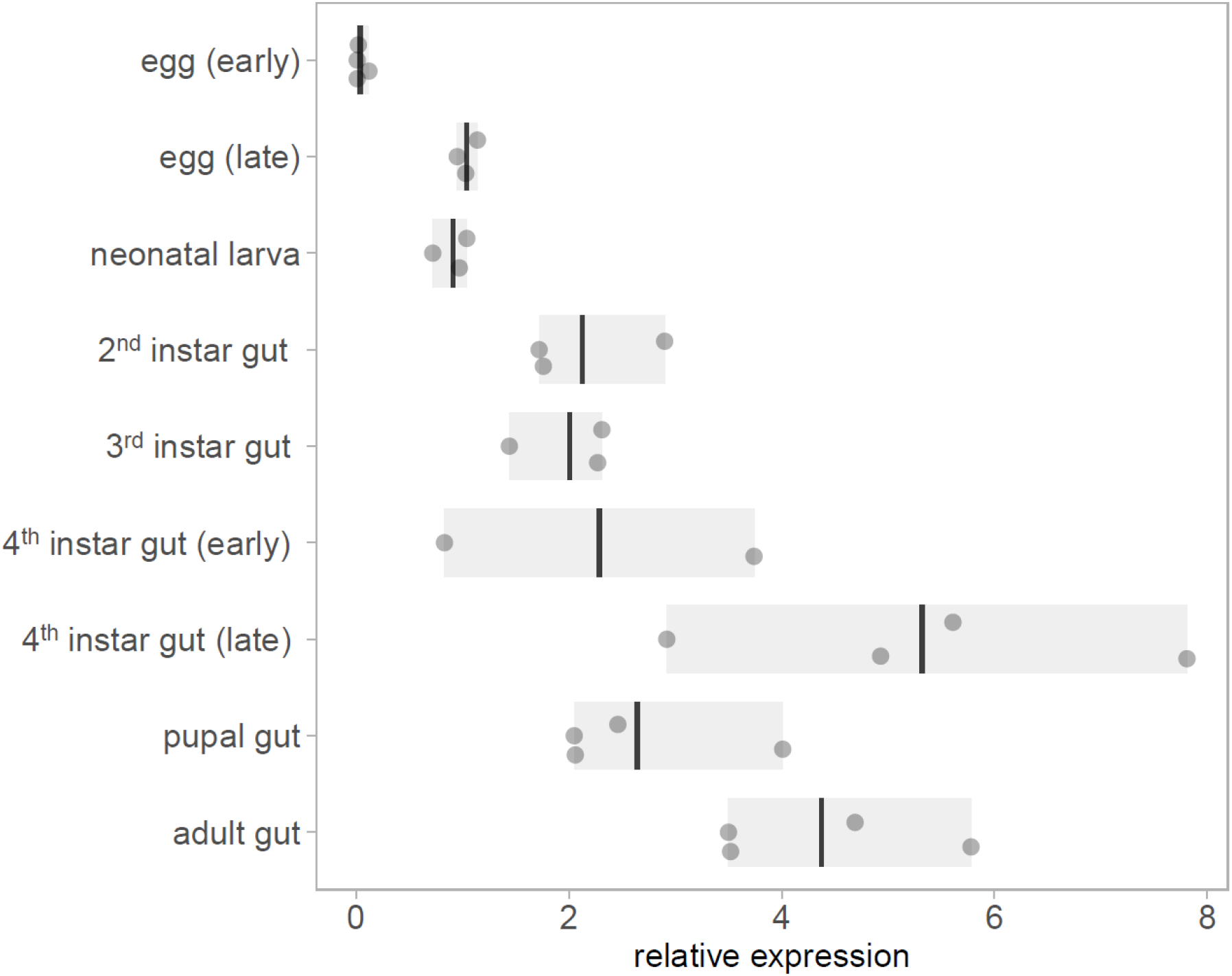
Expression of *mesh* at different CPB developmental stages. Gene expression values are shown relative to early egg sample average. In the eggs and neonatal larval stages, entire organisms were sampled, whereas in later stages, only guts were sampled. Bars show the range of data and lines shown the mean.

### Laboratory feeding trials confirm dsMESH efficiency at different CPB life stages

To test the efficiency of dsMESH in silencing the target gene and its potential as a bioinsecticide we performed three laboratory feeding trials in which we treated CPB at different life stages (Table 1). Firstly, we fed 2^nd^ instar larvae continuously on *in vitro* synthesised dsRNA-sprayed potato foliage and left them to pupate and emerge as adults (trial one, Table 1). We confirmed silencing of *mesh* gene by dsMESH after 4 days of treatment. Compared to dsEGFP treatment, dsMESH reduced *mesh* expression by 71% (p<0.001, Supplementary Dataset 3) and larval survival at that point was 48%, 80%, and 95% for dsMESH, dsEGFP, and water treatment, respectively (Supplementary Figure 4 and Supplementary Dataset 4).

We also tested the effectiveness of dsMESH on 4^th^ instar larvae (trial two, Table 1), which is the final instar before pupation. We recorded adult emergence and inspected plant substrate for beetle carcasses at the end of the trial. Adult emergence rate was 11% in larvae exposed to dsMESH, which is significantly lower compared to more than 75% for dsEGFP and water treatments (p<0.01; Figure 2B and Supplementary Dataset 5). Additionally, in all three emerged adults from the dsMESH treated group we observed darkened deformed elytra (Supplementary Figure 5) and the beetles died within two days after emergence. In contrast, dsEGFP and water treated beetles exhibited normal phenotypes and no adult mortality. From the substrate of dsMESH treated plants, we recovered two larval and six adult carcasses (Supplementary Figure 5) whereas in substrates of dsEGFP and water treated plants we found no carcasses. Trial two thus shows that dsMESH is also effective against 4^th^ instar larvae.

**Figure 2.**
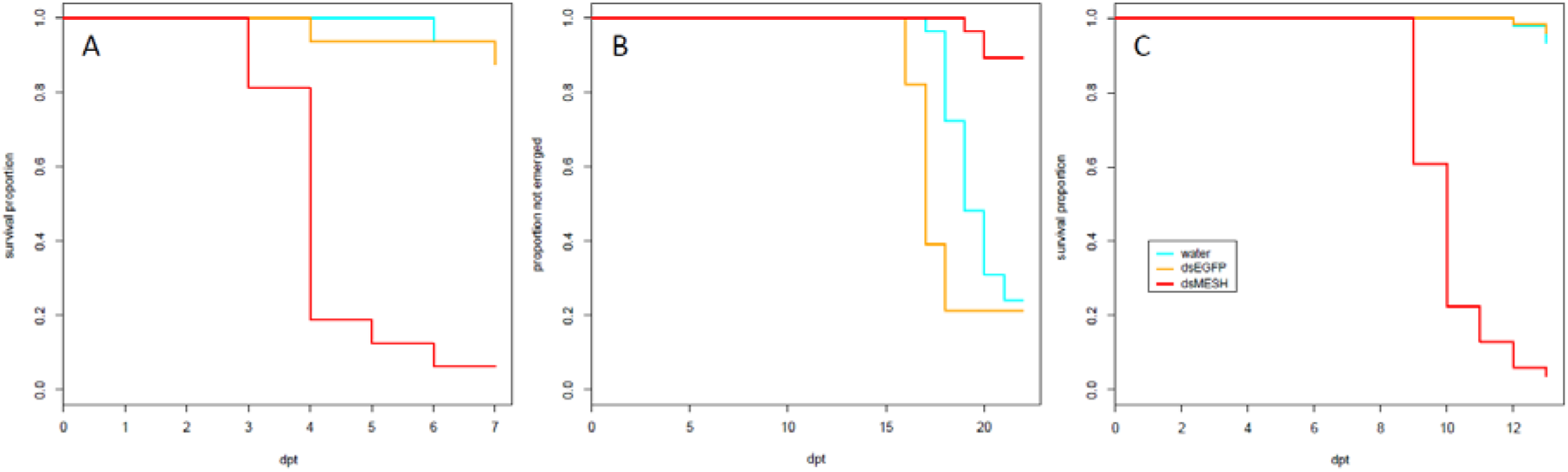
Effectiveness of *in vitro* synthesised dsMESH treatment of different Colorado potato beetle developmental stages. **(A)** Kaplan-Meier survival curves of 2^nd^ instar larvae exposed to discontinuous administration of dsRNA (trial three). Survival was significantly reduced for dsMESH compared to water or dsEGFP treatment (p<0.001). **(B)** Proportion of beetles not emerged as adults after 4^th^ instar larval dsRNA treatment (trial two). Adult emergence was significantly lower for dsMESH treated larvae compared to water or dsEGFP treatment (p<0.01). **(C)** Kaplan-Meier survival curves of larvae hatched from dsRNA-treated eggs (trial five). Survival was significantly reduced for dsMESH compared to water or dsEGFP treatment (p<0.001). dpt – days post treatment.

In trial five, we tested the effectiveness of dsMESH spraying on CPB eggs. We sprayed freshly laid CPB egg masses (Supplementary Figure 6) and transferred 1^st^ instar larvae to untreated potato foliage. Most larvae hatched three days after egg treatment. We observed no difference in larval emergence between dsMESH and control treatments. Massive larval die-off in dsMESH treated group occurred in 6-7 days old larvae (9-10 days post egg treatment, Figure 2C). The survival of dsMESH treated 6 days old larvae was 61%, and a day later only 23%. In comparison, for both dsEGFP and water treated groups, the survival at that time point was 100% (Figure 2C and Supplementary Dataset 8). Only 4% of dsMESH treated larvae survived until 13 dpt, whereas 97% and 95% larvae survived in dsEGFP and water treated groups, respectively (p<0.001; Figure 2C and Supplementary Data 1.9). This trial shows high insecticidal efficiency of dsMESH also when sprayed on CPB eggs.

### The treatment regime does not affect the efficiency of dsRNA

We next compared the effect of the dsRNA administration approach. Contrary to the above-described feeding trials, where larvae were continuously fed with dsRNA-sprayed potato leaves, here we exposed each individual larva (2^nd^ instar) to the same dose of dsRNA by discontinuous administration via treated potato leaf disks (trial three and four, Table 1). We observed similar survival trends as the ones obtained with continuous treatment regime (trial one).

In the first leaf-disk feeding trial (trial three), we observed a substantial reduction in survival 4 dpt, reaching only 18% survival in the case of dsMESH treated larvae compared to more than 90% survival in dsEGFP and water treatments (Figure 2A, Supplementary Dataset 6). In trial four, most substantial survival reduction was observed 5 dpt, where dsMESH treated larval survival rate was 54% compared to 100% survival in both dsGFP and water treatments (Figure 3, Supplementary Dataset 7). In both trials, statistical analysis indicates highly significant survival reduction for dsMESH treatment (p<0.001; Supplementary Data 1.9).

**Figure 3.**
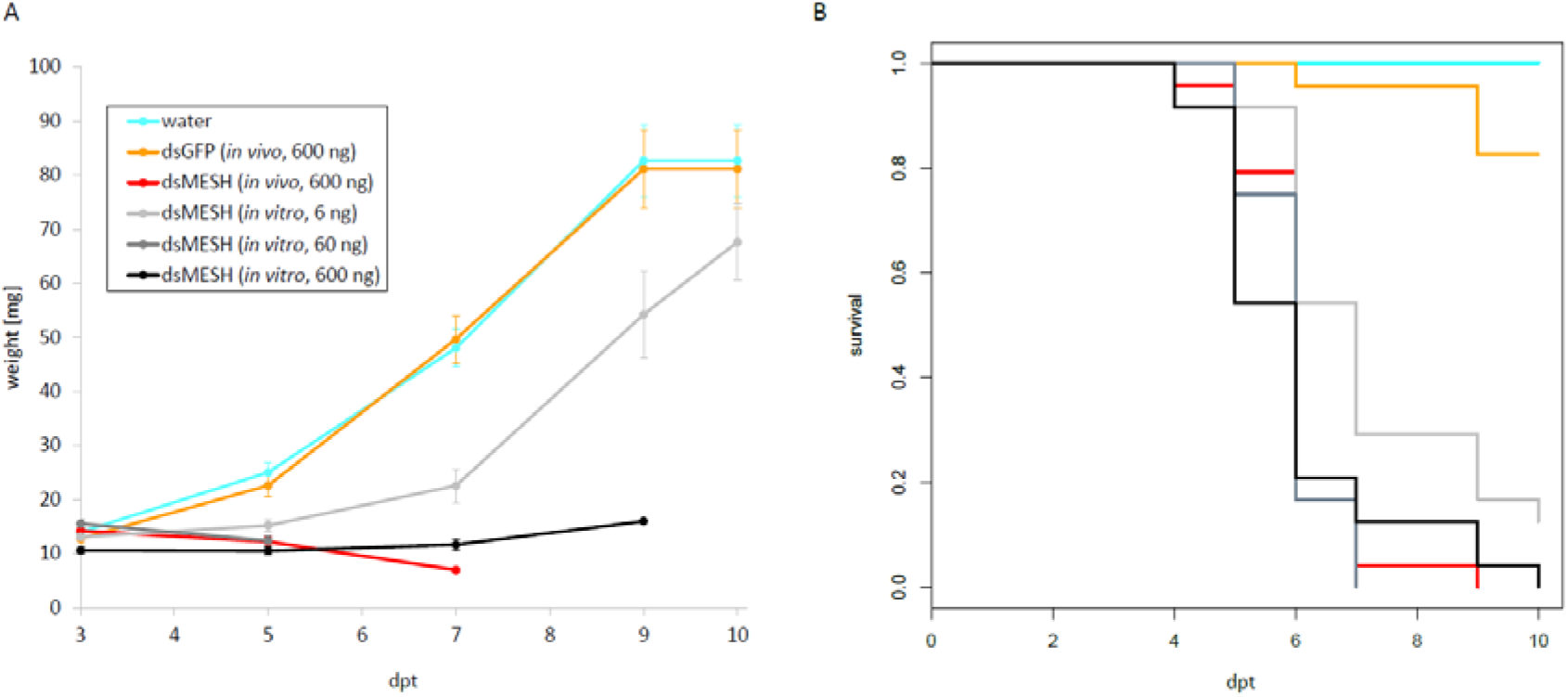
Efficiency of *in vitro* and *in vivo* synthesised dsMESH (trial four). **(A)** Larval weight throughout the trial was significantly reduced when using either *in vitro* or *in vivo* synthesised dsMESH compared to water or dsEGFP treatment. Error bars show standard error of the mean. **(B)** Kaplan-Meier survival curves with survival proportions plotted. Survival was significantly reduced for dsMESH compared control treatments (p<0.001). dpt – days post treatment.

### Comparison of *in vivo* and *in vitro* synthesised dsRNA efficiency

Larval survival analysis and weight measurements in trial four (Figure 3; Supplementary Datasets 7 and 12) show that *in vivo* and *in vitro* synthesised dsMESH are similarly effective (Supplementary Data 1.9). In addition, by testing serial dilutions of *in vitro* synthesised dsMESH, we showed that ingestion of as little as 6 ng of dsMESH caused more than 90% larval mortality (Figure 3B). No significant effect of bacterially produced dsGFP treatment on larval weight or survival was observed (Figure 3).

### The dsMESH treatment against CPB is also efficient in the field

In order to confirm the efficacy of dsMESH as potential insecticide also under environmental conditions we treated potato plants growing in three different fields with *in vitro* synthesised dsMESH. No formulation to increase dsRNA stability or uptake was used to make the results of the field trial more comparable to the laboratory-gained results. Mortality rates for dsMESH treatment after 7 days were significantly higher (F_2, 40_=16; P<0.0001) compared to untreated plants according to ANOVA and were 93%, 84%, and 95%, in the three locations, respectively. GLM analyses showed a significant effect of factor treatment on parameters leaf damage increase (F_2, 41_=34, 8; P<0.0001) and insect mortality rate (F_2,40_=13.2; P<0.0001; Figure 4). Factors experiment and replicate did not significantly affect the observed parameters. Spinosad acted more rapidly than dsMESH, causing on average 98% of larval mortality in just two days, whereas the average mortality rate of dsMESH treatment at that time point was 32% (Supplementary Dataset 13).

**Figure 4.**
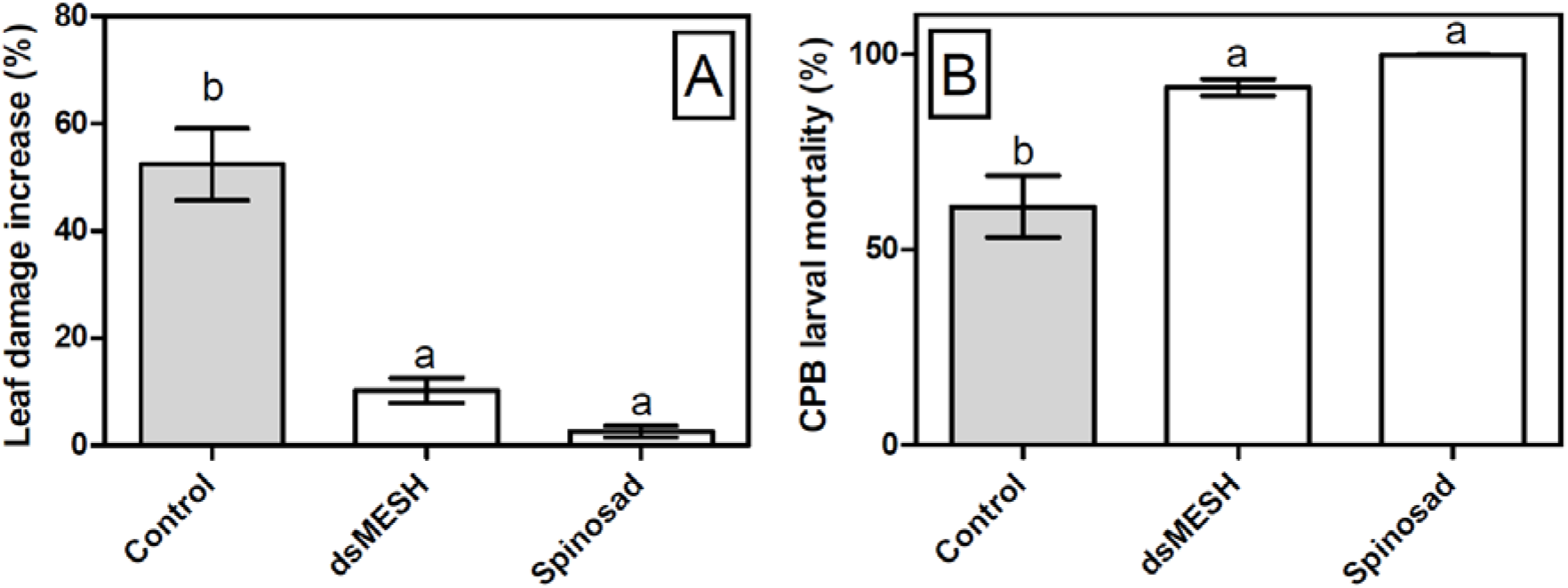
Treatment with dsMESH reduced Colorado potato beetle potato infestation in the field. **(A)** Potato defoliation due to CPB herbivory expressed as leaf damage increase, which was calculated by subtracting data from initial and final leaf damage assessment after seven days. **(B)** CPB larval mortality in the field trial. The data from all three locations were used. Bars not sharing the same lowercase letter are significantly different.

## DISCUSSION

We performed a systematic evaluation of applicability of an RNAi-based insecticide targeting the *mesh* gene (dsMESH) and validated its insecticidal action in CPB. The incentive to use this target gene came from its high expression in CPB gut in most developmental stages and the lethal phenotypes observed in Drosophila knockout mutants (Gramates et al. 2017) and *Tribolium castaneum* RNAi screens (Ulrich et al. 2015). Mesh is a transmembrane protein important for proper organisation of the insect midgut septate junctions and Drosophila *mesh* mutants show an impaired barrier function of the midgut (Izumi et al. 2012). Silencing *mesh* by RNAi in Drosophila adults, however, does not impair gut integrity but increases gut bacterial load by regulating dual oxidase expression (Xiao et al. 2017).

Mesh was first identified as an effective RNAi pesticide target in western corn rootworm, *Diabrotica virgifera virgifera*, another coleopteran pest closely related to CPB (Hu et al. 2016). Our CPB feeding trials with *in vitro* synthesized dsMESH consistently showed high mortality rates in larvae with effective dose in the ng range, similarly as reported for corn rootworm by Hu et al. (2016). In our first feeding trials we used 2^nd^ instar larvae because the first two CPB instars were described as most susceptible to RNAi (Guo et al. 2015). In addition, we showed that dsMESH treatment is effective against 4^th^ instar larvae and CPB eggs. Surprisingly, reports of insect egg treatment by spraying or soaking in dsRNA are rare and have different outcomes. Soaking Asian corn borer *(Ostrinia furnalalis)* eggs in pesticidal dsRNA solutions caused reduced hatching (Wang et al. 2011). On the contrary, in the corn earworm, *Helicoverpa zea*, soaking eggs in dsRNA as well as larval feeding delivery had no effect, whereas injecting eggs with same dsRNA induced RNAi and reduced egg hatching rate (Wang et al. 2018). In our trial, spraying eggs with dsMESH did not affect egg hatching although we showed that *mesh* is expressed in eggs. The larval die-off six to seven days after emergence from dsMESH treated eggs leads to suggest that dsRNA was mostly taken up by neonatal larvae while feeding on eggshells.

The activity of dsRNA obtained in *in vitro* tests or laboratory feeding experiments might not reflect that on the field. Thus, we decided to validate our laboratory-based trial results in a field trial comparing dsMESH efficiency to that of spinosad. Spinosad was used as a positive control as a) it is highly effective against CPB, b) it is a bioinsecticide and can thus be used also in organic farming, c) it is an insecticide registered for control of CPB in Slovenia and d) it is an alternative to conventional chemical insecticides (e.g. thiacloprid, beta-cyfluthrin), which can be ecotoxicologically problematic, and for which we are trying to find alternatives for. The observed field mortality was slightly lower compared to laboratory trials, which is reasonable, as larval treatment on the field was not as controlled and uniform as in the laboratory. In addition, reduced dsRNA stability in the field is expected due to direct sunlight exposure and lack of formulation to improve dsRNA stability (Cagliari et al. 2019). Compared to the wide-spectrum insecticide spinosad (Kirst 2010), dsMESH has an inherent lag phase in observed mortality, which can be attributed to its mode of action. The toxicity of dsRNA depends on target protein’s halflife (Scott et al. 2013) therefore, we expected to observe lethal effects after a few days. Despite its slower activity, the final mortality and leaf damage caused by dsMESH treatment in the field trial were not statistically different to that of spinosad.

In our experiments we used *in vitro* synthesised dsRNA, however, for large scale field application applying crude extract of *E. coli* producing dsMESH might be a good option to reduce the costs. Our laboratory feeding trials showed that *in vivo* produced dsMESH and the dsMESH synthesized *in vitro* are similarly effective. Because the dsMESH amounts in bacterial extracts was approximated from the gel, a more accurate comparison is not possible. Despite the potential advantages of applying dsRNA as a crude bacterial extract, the approach has also additional risks such the presence of synthetic DNA elements and the possibility of having a GMO status assigned (Fletcher et al. 2020).

## CONCLUSIONS

Although plant-incorporated protectants (transgenic plants) are the most cost-effective way of using RNAi-based pesticide technology, their public acceptance might prove challenging, at least in the European Union. Other possibilities were envisioned, such as transformed insect symbionts (Whitten et al. 2016) or viruses expressing pesticidal RNA molecules (Taning et al. 2018), albeit again using genetically modified organisms. As an alternative, non-transformative strategies of dsRNAs application, i.e. spray-induced gene silencing, are being investigated and CPB is the first agricultural pest for which this technology might be commercialized (Cagliari et al. 2019).

We have shown in laboratory trials as well as in the field that spraying with insecticidal dsRNA is a highly efficient strategy for managing CPB. We are planning to test dsMESH in a larger field trial using standard agricultural spraying equipment and against a range of other insecticides. For RNAi-recalcitrant agricultural pests, future research will have to focus on formulations to improve dsRNA stability and cellular uptake. Apart from efficiency, further research is needed on biosafety implications of this new pest management strategy. This includes investigating possible impact of dsRNA on human health and the environment (Rodrigues and Petrick 2020). For sustainable use of this technology in agriculture, integrated pest management strategies will have to be employed to delay the development of pest resistance to dsRNAs (Khajuria et al. 2018).

## AUTHOR CONTRIBUTIONS

MP provided the initial concept and design of the study, performed gene expression analysis, laboratory-based feeding trials, contributed to execution and evaluation of the field trial and wrote the manuscript. AC and RF established the bacterial production of dsRNA. RF also helped with execution of the feeding trial four. JR designed and led the execution of the field trial. KG contributed to study design, data interpretation and manuscript drafting. All authors contributed to manuscript revision, read and approved the submitted version.

## ACKNOWLEDGMENTS

The authors thank Špela Prijatelj Novak, Tjaša Lukan, Lidija Matičič and Eva Praprotnik for technical support and prof. Guy Smagghe for supplying CPB eggs from his lab colony. The *E. coli* HT115 (DE3) strain was provided by the CGC, which is funded by NIH Office of Research Infrastructure Programs (P40 OD010440).

## REFERENCES

Albright VC, Wong CR, Hellmich RL, Coats JR (2017) Dissipation of double-stranded RNA in aquatic microcosms. Environ Toxicol Chem 36:1249–1253. doi: 10.1002/etc.3648

Alyokhin A, Baker M, Mota-Sanchez D, et al (2008) Colorado Potato Beetle Resistance to Insecticides. Am J Potato Res 85:395–413. doi: 10.1007/s12230-008-9052-0

Baebler Š, Svalina M, Petek M, et al (2017) quantGenius: implementation of a decision support system for qPCR-based gene quantification. BMC Bioinformatics 18:276. doi: 10.1186/s12859-017-1688-7

Baum J a, Bogaert T, Clinton W, et al (2007) Control of coleopteran insect pests through RNA interference. Nat Biotechnol 25:1322–1326. doi: 10.1038/nbt1359

Cagliari D, Dias NP, Galdeano DM, et al (2019) Management of Pest Insects and Plant Diseases by Non-Transformative RNAi. Front Plant Sci 10:1319. doi: 10.3389/fpls.2019.01319

Cappelle K, de Oliveira CFR, Van Eynde B, et al (2016) The involvement of clathrin-mediated endocytosis and two Sid-1-like transmembrane proteins in double-stranded RNA uptake in the Colorado potato beetle midgut. Insect Mol Biol 25:315–323. doi: 10.1111/imb.12222

Christiaens O, Dzhambazova T, Kostov K, et al (2018) Literature review of baseline information on RNAi to support the environmental risk assessment of RNAi-based GM plants. EFSA Support Publ 15:1424E. doi: 10.2903/sp.efsa.2018.EN-1424

Dubelman S, Fischer J, Zapata F, et al (2014) Environmental Fate of Double-Stranded RNA in Agricultural Soils. PLoS One 9:e93155. doi: 10.1371/journal.pone.0093155

EPPO (2008) Efficacy evaluation of insecticides, Leptinotarsa decemlineata, PP1/012(4)

Fletcher SJ, Reeves PT, Hoang BT, et al (2020) A Perspective on RNAi-Based Biopesticides. Front Plant Sci 11:51. doi: 10.3389/fpls.2020.00051

Fu K-Y, Li Q, Zhou L-T, et al (2016) Knockdown of juvenile hormone acid methyl transferase severely affects the performance of *Leptinotarsa decemlineata* (Say) larvae and adults. Pest Manag Sci 72:1231–1241. doi: 10.1002/ps.4103

Fu K-Y, Lü F-G, Guo W-C, et al (2015) Characterization and functional study of a putative juvenile hormone diol kinase in the Colorado potato beetle *Leptinotarsa decemlineata* (Say). Arch Insect Biochem Physiol 90:154–167. doi: 10.1002/arch.21251

Gramates LS, Marygold SJ, Santos G dos, et al (2017) FlyBase at 25: looking to the future. Nucleic Acids Res 45:D663–D671. doi: 10.1093/nar/gkw1016

Guo W-C, Liu X-P, Fu K-Y, et al (2016) Nuclear receptor ecdysone-induced protein 75 is required for larval-pupal metamorphosis in the Colorado potato beetle *Leptinotarsa decemlineata* (Say). Insect Mol Biol 25:44–57. doi: 10.1111/imb.12197

Guo W, Bai C, Wang Z, et al (2018) Double-Stranded RNAs High-Efficiently Protect Transgenic Potato from *Leptinotarsa decemlineata* by Disrupting Juvenile Hormone Biosynthesis. J Agric Food Chem 66:11990–11999. doi: 10.1021/acs.jafc.8b0391

Guo Z, Kang S, Zhu X, et al (2015) The novel ABC transporter ABCH1 is a potential target for RNAi-based insect pest control and resistance management. Sci Rep 5:13728. doi: 10.1038/srep13728

Horn T, Boutros M (2010) E-RNAi: A web application for the multi-species design of RNAi reagents-2010 update. Nucleic Acids Res 38:332–339. doi: 10.1093/nar/gkq317

Hu X, Richtman NM, Zhao J-Z, et al (2016) Discovery of midgut genes for the RNA interference control of corn rootworm. Sci Rep 6:30542. doi: 10.1038/srep30542

Hu X, Steimel JP, Kapka-Kitzman DM, et al (2019) Molecular characterization of the insecticidal activity of double-stranded RNA targeting the smooth septate junction of western corn rootworm (*Diabrotica virgifera virgifera*). PLoS One 14:e0210491. doi: 10.1371/journal.pone.0210491

Izumi Y, Yanagihashi Y, Furuse M (2012) A novel protein complex, Mesh-Ssk, is required for septate junction formation in the Drosophila midgut. J Cell Sci 125:4923–4933. doi: 10.1242/jcs.112243

Joga MR, Zotti MJ, Smagghe G, Christiaens O (2016) RNAi Efficiency, Systemic Properties, and Novel Delivery Methods for Pest Insect Control: What We Know So Far. Front Physiol 7:553. doi: 10.3389/fphys.2016.00553

Khajuria C, Ivashuta S, Wiggins E, et al (2018) Development and characterization of the first dsRNA-resistant insect population from western corn rootworm, *Diabrotica virgifera virgifera* LeConte. PLoS One 13:e0197059. doi: 10.1371/journal.pone.0197059

Kirst HA (2010) The spinosyn family of insecticides: realizing the potential of natural products research. J Antibiot (Tokyo) 63:101–111. doi: 10.1038/ja.2010.5

Lü F-G, Fu K-Y, Guo W-C, Li G-Q (2015) Characterization of two juvenile hormone epoxide hydrolases by RNA interference in the Colorado potato beetle. Gene 570:264–271. doi: 10.1016/j.gene.2015.06.032

Meng Q-W, Liu X-P, Lü F-G, et al (2015) Involvement of a putative allatostatin in regulation of juvenile hormone titer and the larval development in *Leptinotarsa decemlineata* (Say). Gene 554:105–113. doi: 10.1016/j.gene.2014.10.033

Meng QW, Xu QY, Deng P, et al (2018) Transcriptional response of Methoprene-tolerant (Met) gene to three insect growth disruptors in *Leptinotarsa decemlineata* (Say). J Asia Pac Entomol 21:466–473. doi: 10.1016/j.aspen.2018.02.011

Petek M, Rotter A, Kogovšek P, et al (2014) *Potato virus Y* infection hinders potato defence response and renders plants more vulnerable to Colorado potato beetle attack. Mol Ecol 23:5378–5391. doi: 10.1111/mec.12932

Petek M, Turnšek N, Gašparič MB, et al (2012) A complex of genes involved in adaptation of *Leptinotarsa decemlineata* larvae to induced potato defense. Arch Insect Biochem Physiol 79:153–181. doi: 10.1002/arch.21017

Rice P, Longden I, Bleasby A (2000) EMBOSS: The European Molecular Biology Open Software Suite. Trends Genet 16:276–277. doi: 10.1016/S0168-9525(00)02024-2

Rodrigues TB, Petrick JS (2020). Safety Considerations for Humans and Other Vertebrates Regarding Agricultural Uses of Externally Applied RNA Molecules. Front Plant Sci 11:407. doi: 10.3389/fpls.2020.00407

Schoville SD, Chen YH, Andersson MN, et al (2018) A model species for agricultural pest genomics: the genome of the Colorado potato beetle, *Leptinotarsa decemlineata* (Coleoptera: Chrysomelidae). Sci Rep 8:1931. doi: 10.1038/s41598-018-20154-1

Scott JG, Michel K, Bartholomay LC, et al (2013) Towards the elements of successful insect RNAi. J Insect Physiol 59:1212–1221. doi: 10.1016/j.jinsphys.2013.08.014

Shi J-F, Fu J, Mu L-L, et al (2016a) Two *Leptinotarsa* uridine diphosphate N-acetylglucosamine pyrophosphorylases are specialized for chitin synthesis in larval epidermal cuticle and midgut peritrophic matrix. Insect Biochem Mol Biol 68:1–12. doi: 10.1016/j.ibmb.2015.11.005

Shi J-F, Mu L-L, Chen X, et al (2016b) RNA interference of chitin synthase genes inhibits chitin biosynthesis and affects larval performance in *Leptinotarsa decemlineata* (Say). Int J Biol Sci 12:1319–1331. doi: 10.7150/ijbs.14464

Shi X-Q, Guo W-C, Wan P-J, et al (2013) Validation of reference genes for expression analysis by quantitative real-time PCR in *Leptinotarsa decemlineata* (Say). BMC Res Notes 6:93. doi: 10.1186/1756-0500-6-93

Taning CNT, Christiaens O, Li X, et al (2018) Engineered Flock House Virus for Targeted Gene Suppression Through RNAi in Fruit Flies *(Drosophila melanogaster) in Vitro* and *in Vivo*. Front Physiol 9:805. doi: 10.3389/fphys.2018.00805

Therneau TM, Grambsch PM (2000) Modeling Survival Data: Extending the Cox Model. New York: Springer New York.

Ulrich J, Dao VA, Majumdar U, et al (2015) Large scale RNAi screen in Tribolium reveals novel target genes for pest control and the proteasome as prime target. BMC Genomics 16:674. doi: 10.1186/s12864-015-1880-y

Wan P, Fu K, Lü F, et al (2014) A putative Δ1-pyrroline-5-carboxylate synthetase involved in the biosynthesis of proline and arginine in *Leptinotarsa decemlineata*. J Insect Physiol 71:105–113. doi: 10.1016/j.jinsphys.2014.10.009

Wang J, Gu L, Knipple DC (2018) Evaluation of some potential target genes and methods for RNAi-mediated pest control of the corn earworm *Helicoverpa zea*. Pestic Biochem Physiol 149:67–72. doi: 10.1016/j.pestbp.2018.05.012

Wang Y, Zhang H, Li H, Miao X (2011) Second-Generation Sequencing Supply an Effective Way to Screen RNAi Targets in Large Scale for Potential Application in Pest Insect Control. PLoS One 6:e18644. doi: 10.1371/journal.pone.0018644

Whitten MMA, Facey PD, Del Sol R, et al (2016) Symbiont-mediated RNA interference in insects. Proc R Soc B Biol Sci 283:20160042. doi: 10.1098/rspb.2016.0042

Xiao X, Yang L, Pang X, et al (2017) A Mesh–Duox pathway regulates homeostasis in the insect gut. Nat Microbiol 2:17020. doi: 10.1038/nmicrobiol.2017.20

Xu Q-Y, Meng Q-W, Deng P, et al (2018) *Leptinotarsa* hormone receptor 4 (HR4) tunes ecdysteroidogenesis and mediates 20-hydroxyecdysone signaling during larval-pupal metamorphosis. Insect Biochem Mol Biol 94:50–60. doi: 10.1016/j.ibmb.2017.09.012

Zhou L-T, Jia S, Wan P-J, et al (2013) RNA interference of a putative S-adenosyl-L-homocysteine hydrolase gene affects larval performance in *Leptinotarsa decemlineata* (Say). J Insect Physiol 59:1049–1056. doi: 10.1016/j.jinsphys.2013.08.002

Zhu F, Xu J, Palli R, et al (2011) Ingested RNA interference for managing the populations of the Colorado potato beetle, *Leptinotarsa decemlineata*. Pest Manag Sci 67:175–182. doi: 10.1002/ps.2048

Zhu T-T, Meng Q-W, Guo W-C, Li G-Q (2015) RNA interference suppression of the receptor tyrosine kinase Torso gene impaired pupation and adult emergence in *Leptinotarsa decemlineata*. J Insect Physiol 83:53–64. doi: 10.1016/j.jinsphys.2015.10.005

