## Supplementary Data, Figures and Tables for "Validating the potential of double-stranded RNA targeting Colorado potato beetle *mesh* gene in laboratory and field trials"

### *Supplementary Material*

**Table of contents**

**1 Supplementary Data**
1.1 Correction of mesh gene model and dsMESH design
1.2 dsRNA sequences used in the trials
1.3 Evaluation of *in vivo* dsRNA production
1.4 Feeding trial one: continuous treatment, 2nd instar larvae
1.5 Feeding trial two: continuous treatment, 4th instar larvae
1.6 Feeding trial three: discontinuous leaf disk treatment with dsRNA, 2nd instar larvae
1.7 Feeding trial four: discontinuous leaf disk treatment with *in vitro* and *in vivo* synthesised dsRNA, 2nd instar larvae
1.8 Feeding trial five: egg treatment
1.9 Survival analysis statistics for laboratory feeding trials

**2 Supplementary Figures**
Supplementary Figure 1. Correction of mesh gene model
Supplementary Figure 2. Estimation of *in vivo* synthesised dsRNA amounts
Supplementary Figure 3. Expression of *mesh* gene in Colorado potato beetle body parts
Supplementary Figure 4. Experimental measurements of continued feeding trial of 2nd instar larvae with *in vitro* synthesised dsRNA (trial one)
Supplementary Figure 5. Colorado potato beetle phenotypes observed in adults emerged from 4th instar larvae continuously exposed to dsMESH (trial two)
Supplementary Figure 6. Spraying of Colorado potato beetle eggs (trial five)

**3 Supplementary Tables**
Supplementary Table 1 List of qPCR assays used in this study and their properties according to MIQE guidelines

**Supplementary Datasets** 1-13 are available as a separate Excel file.

### 1 SUPPLEMENTARY DATA

#### 1.1 Correction of *mesh* gene model and dsMESH design

To correct the i5k genome *mesh* model (genome version 0.5.3), we used the exon junction evidence of the mapped transcriptome assembly contig 1:CUFF.34151.1 that corresponded best to mapped Illumina RNA-Seq reads (Supplementary Figure 1A). To obtain the longest possible transcript, another exon supported by Illumina RNA-Seq reads originating from augustus_masked-Scaffold721-abinit-gene-8.2-mRNA-1 model was added to the manually corrected model (exon11). The final corrected *mesh* model consists of 13 exons (sequences listed below) and the translated protein matches best the Colorado potato beetle (CPB) protein *mesh* isoform X3 predicted by NCBI genome annotation pipeline although missing the exon11 sequence (Supplementary Figure 1B). Due to uncertainty in exon structure at 3'-end we decided to exclude exon11-13 from dsRNA design.

```
>LDEC006484_manually_corrected_exon1
TTAATACAGACTCAGATAACAAATTTTATCTTGTTACAGAGCACACTTGTGAGAAGGCAATCGAGTAACGAACCTCATACATAAAAGTACACGTATGGTGATAGAAA
ACAATAAATTATCAAACTGAGGCTCTGTGCCAAGTTTTGGAAAATGTACGTCAAGTGGAAATTGGTTTTGACTCTAGTACTGTGTGTTGGTGTCGTGATTGGGGAAG
ATATTTCAACAGATATAATCCCCTTGCCACAAGACAATACAGCAGATGTGGAAATAGTGGCTACAGAAACTAGGTCAAATTCAGGATCTGAAGCTGCTGTAGAAGAG
CCCACCAATAACACTGGGCCTTCAGATACTACAAACCCAAATTATG
>LDEC006484_manually_corrected_exon2
CTGTAGTGCCCCCTGTAGTTTCTGATAGCAAACCTGCAATAACAAATAGTGAAACAAATGACACTAACAATGATGTTGTAATGTTGAGTCCTACGAAATTTCTCATG
AAAAAGGGACGATCTGGACGTTTATTGGAATATCCGACTGATTATG
>LDEC006484_manually_corrected_exon3
ATCCTATGACTTCCCATATTGCTCCACCGGATAGTGATCAGAGAGGATATTCAGGAGTACCATATGTCTTGACGGAAACCAGACTGCAACAGATCCGCCAAAATTTC
ATGTATCCCTACTACAACAGAGGCGGTAATGCAGATGACGAAGGAGACTACCAGAAAGAAATTCAATCATCTATTCCGCAAGTGTACAAGAACCTCAACTTCCAACT
CCCTTTCTTCGGATTTCGATTCAATTACACGAGGGTCTCCTTGAACGGTTATTTGGAATTCAGCGATCCTCCTCCAAATTACGACTATCCTTTGGTCTTTCCAGTAA
AGGAATGGCCTAAAAAGAACGATCCTTCTTTCATCGGTATCTTTTTCAGTAAATGTAGAATCGGTAACCTGAGGGACGGAGATATTGATCAAAGAGACCCTGGAGTG
TACTTTAGGATGGAAAGGGATCTCAGAAATAGGCAGGACAGGATGGGAGTGGAGATCAGAGAACGACTGAAATGGGATATAAGGGAAGGGGTGATAGGGTCAGAAAC
ATTCAATCCCAAACACGCCATTATCGTCACATGGAAAAATATCTCTTTCAATGGAGGTTTTGGCAATGCTCTCTACCAG
>LDEC006484_manually_corrected_exon4
ACTAACACTTTCCAAATGATCCTCGCCACTGATGAAGTTTTCACCTACGCCATGTTCAACTACTTGAATCTTGACTGGACCACCCACACTGAAGCGGGAGGCGACAC
AAGAAAAGGAGAAGGAGGAGTTCCCGCTTTTGTGGGATTCAACGCTGGAAACGGTACTAGAAGTTTTGAATACAAACCATACAGTCAAGAATCTGTTATTCGAGATC
TCACACAAACTGGTTTCGCTAATGGTTTCAAAGGAAGGCACATTTTCCGAATCGACGAAAATATCCTAACTGGAACATGCAATAAAGATATAG
>LDEC006484_manually_corrected_exon5
ATGGTGCTAATCTACCGTTAATGATATCTCCAGAAAGTGGAAATATGCTGGGTGGAACAATAGTGAATATAACAGGACCTTGTTTCGGCCTAGACGACCAAGTTAAA
TGCAAATTTGATGTAGCCGATGAAATAAATGGCGTCGTTATAGATAAAAACAGGGCTATATGCATCCAACCTAGACTGTATGCCGAAGGATGGGTGAATTTACAAAT
AGCCATAGGGGCTGGGGTATACAAATGGAAGGGAAAATATTATGTCG
>LDEC006484_manually_corrected_exon6
AATCCCCCGCAGCGGCATCTCAAAAAATCTACTTCAAGGACATGAAGGTTCATGAAAAATCGCCTAGTGAAATAAGAATAACTTGGGAAAAATACAACTTGACCACT
AACGAAAACGCTAACATTCGCATCTCCTTATGGGGTTACAGAGAAACAACAATAAGGCCAACGTTCGTTTACATCACTGATATCGCAGACAGTCTCCAAAATACTGG
AGAGTATACCATCGTACCGTCCCAATATAGAACAAAAGTTAATGAGTTTCTCACGGATATCAAATTTGGTTTCTTGCAAATTAACTTGACTGAATCTATCAAA
>LDEC006484_manually_corrected_exon7
GTGAACACTTATACATCTGTACAACGATCAGTGGAAATAGTTCCTGTTGTATGGAGTCGACCCATTCCCCTAGGATGGTACTTTCAGTTCCAATGGGAAAATATGTA
TGGACGAAGCTGGCCCAAAGCACTCTGCGATGACTGGCTAAGAACAGACAGATACCTGAAAAACTTTGCTCATGAGTTACCTCAATGCCCTTGCACTGTAGAACAGG
CTTTGGCAGACAAAGGGAGGTATATGCCCGACTTTGATTGCGACAAGGACTCAAATCCCGTATGCTACTACAATAACCAAGCTCTGCACTGTGTGAAAACAGGATCA
CCAAC
>LDEC006484_manually_corrected_exon8
GTTGGAGGGATCAGAACAGCAGTGCTGCTATGACAAAAACGGGTATCTCATGTTATCATACGATCAGCAGTGGGGTTCAAGTCCACGGCGTTGCCACAATCTGGGAA
AAATGCCCTACAACGAAGCAACAAAAGTTCCAACCTTATCGCAATGGTTCAACGATATGGTACCGAAGTATCTTTGCTGTTTGTGGCAGGAAGAACAGGCGGTGGGT
TGCGAAACGCTGAGATTCGAAAGAAGACCAACTCAGGACTGTGTCGCGTACCAAGCTCCAGGGATTGCTGGGATTTACGGAGATCCCCACGTCATCACTTTCGATGA
CGTCGAGTACACCTTCAACGGGAAAGGAGAGTTTGCTCTTGTGAAATCTGTGACACAAACTGACAACTTGGAGGTGCAAGGCAGATTTGAGCAAATGGACCCTAACG
CCTACGGAGAAGTACGTGCAACACAACTGACTTCAATTGTGGCAAGGGGAAACAACACCATAGCAGTGGAGGTCAGAAGGAGGCCCTTGGATGCTAGGTGGAGGTAT
AGGCTGGATGTCATTGCTGATAATAGGAAGTTGTTCTTCGACAGACCCTCTTTGAAATTCCAACATTTCCAAG
>LDEC006484_manually_corrected_exon9
GAGTGACTATTTATACACCTACTTATATCCTCAATCAGTCTGAAGTCATCATTATGTTTGATAACGGAGCAGGAGTTCAAGTAATGGATAACCAGGGATTCATGACC
GCGAGGGTGTATCTTCCTTGGTCATTCATC
>LDEC006484_manually_corrected_exon10
AACAAAACTGTTGGTCTCTTTGGCAACTGGAGTTTCAATAAGGAAGATGACTTCACTCTTCCTGATGAGTCGAAGGCTGCCGTCGTGGGTAATATCAATGATATGGA
AAGGGTCTACAACGATTTTGGTTCCAAATGGATGGTGGACGACGTACTAGATCCGAAAAGAGGTAGATCCCTATTTTTCAGAGAATTCGGCAGATCATCGGCAACGT
ACAACAACAAAACTTTCAAACCGCAGTTCCTTATGTTACCTGAGGACATAATACCCGCAAACAGGTCGATACAGATACAGAGAACTTACGACATTTGTAGCACAAAA
ATGTACGAATGCTACTACGATTATGCCATGACGCTCAACAGAGATCTTGCCCATTTTACTCAGAATTATAAAGCAACCATATATCAACTCAAAGAAACGACGAGGCA
GAAGGTTGTTTCTTGCGGAGTTCTGGAAACACCGCGATTCGGTAGGAAGAGTACTTTTCTTTTTATACCAGGAACCAAAGTCACTTACGAGTGCAATCAAGACTTCG
TATTGGTGGGAGATCCCAGAAGAGAATGTTTGGCAGATGGCACATGGAATGCTCCTGAATATGGCTACACCGAATGTTTAC
>LDEC006484_manually_corrected_exon11
GTCAACAAGAATATTCTTCCCGAACCGCCATGATCACTTGGTCCATAATCCTTGCAGTTCTTATACCATTAATCCTATTGATACTTTTCGCGGGATACAAAGTGTAT
CAAAAAATCAAAGGCGACTCATGGGAGGACAACGATTCAACTAAACCTAAAAAATTGCAAGCCTTCAATCGGGCACTCAGTCCATACCAAGATGAAGAAGACGACGA
TGATGACGACTATGTTCCCAATCCCAGTGACAAGTCAGAAA
>LDEC006484_manually_corrected_exon12
GTCAACAAGAATATTCTCAGCGCCAATCAGCCATTGCCTCTGGAGCCGTTCTCGCAATAATTATTCCACTAGTTTTATTATTTGTATATCTGGCTTATATGTTCCTC
AAGAAGAAACAGAAAGAACGAGACGAAGAAAATTTACAAACGCAAGCGTACGAGCAACAGAAAAG
>LDEC006484_manually_corrected_exon13
```

ACAAGCTCAGGAAGCTGCTGCTAGAAAATTAACTGCTGCCGAAGAGTACAACTCAGATGAAGACGATAATAACAGCAACGTTACGAGCACAGCAAAAGAAACAACAG
TGTATTAGTATTAGATATGGTAACACTTTTACGATATTATTATTTTGGTTGGCATACTTCTCAGACAAACAGTCCAAATTATTATTTGCAACAAGAACTATTTGAAT
TTCGAAGTTTCCTTTGAAGATCCCAAGTTGCTTATTATCCTAAATTTTTGCAAAATGTCAAAATCGTACTATTCCATCAGTTGTATATTTTATTTTTCTGAGATAGT
CAGGCCAACAGGTATCTAAAGGAGTGAATAACAGATTTTTCATATTTTCCAGTTTTATCTGATTCAATAATGCACTATGGGTTTTTTCTGTTAATGGTTTTTATAAC
ACCTCTGTTGCAAAATACATATTAGGTAATATAAATATACGTAATTAAGTCGTAATTAAGTATTATAACAAAATGGCGTTCTCTATTATTTAAATTGTAATTAGATA
TATATTTTTTATATTCATAATCACTTTTGCATGTAAAGTAGGAGTCCAATTTTCTGTACTTCAACCATCACTATCGCTACTTACCCCGATCACTTTTTGGATGCACA
TTTCATGCACATTTTTGCACATTGTTTGCACAGTAAGTTCACTTTGCAAACAATGTGCGAAGACTATGCGGGAGGACATGTTTATGCAGTAATTTTGTATATATGTG
AGTAGGTATATACCTACTATTTTCATCATACTCTGGTCCTACTTTTCTTATTTACAGTGATAAGATGATGTTTGAAATCTTGAATTAGGTACAATGAATTCATTTTT
GTCAGTTATTTGAAACATTTTCGGATATTTTAATAAGACTGTGTTTTATCAAAGTTCAGTTTGATTGTCCAAATACTAGTAGATCATTAGAGTCCTACTGTAGTTTT
AGCATTTTAATAAAGTAATTCAATATTAATATACAAGTAATCTTGCACTTCTGCTAACTCAAAATTTTGCAAAATTATTCACTCGAATCAGTATTCACACGTTTCTC
TTTCAGAAATAACCAAAATATTTCACCCCATACACAAATTTCACTAAACCTAATGAAAATATATATTCTCCAATAAATTTTTTGAAAACAAGATG

>LDEC006484_manually_corrected_mRNA
TTAATACAGACTCAGATAACAAATTTTATCTTGTTACAGAGCACACTTGTGAGAAGGCAATCGAGTAACGAACCTCATACATAAAAGTACACGTATGGTGATAGAAA
ACAATAAATTATCAAACTGAGGCTCTGTGCCAAGTTTTGGAAAATGTACGTCAAGTGGAAATTGGTTTTGACTCTAGTACTGTGTGTTGGTGTCGTGATTGGGGAAG
ATATTTCAACAGATATAATCCCCTTGCCACAAGACAATACAGCAGATGTGGAAATAGTGGCTACAGAAACTAGGTCAAATTCAGGATCTGAAGCTGCTGTAGAAGAG
CCCACCAATAACACTGGGCCTTCAGATACTACAAACCCAAATTATGCTGTAGTGCCCCCTGTAGTTTCTGATAGCAAACCTGCAATAACAAATAGTGAAACAAATGA
CACTAACAATGATGTTGTAATGTTGAGTCCTACGAAATTTCTCATGAAAAAGGGACGATCTGGACGTTTATTGGAATATCCGACTGATTATGATCCTATGACTTCCC
ATATTGCTCCACCGGATAGTGATCAGAGAGGATATTCAGGAGTACCATATGTCTTGACGGAAACCAGACTGCAACAGATCCGCCAAAATTTCATGTATCCCTACTAC
AACAGAGGCGGTAATGCAGATGACGAAGGAGACTACCAGAAAGAAATTCAATCATCTATTCCGCAAGTGTACAAGAACCTCAACTTCCAACTCCCTTTCTTCGGATT
TCGATTCAATTACACGAGGGTCTCCTTGAACGGTTATTTGGAATTCAGCGATCCTCCTCCAAATTACGACTATCCTTTGGTCTTTCCAGTAAAGGAATGGCCTAAAA
AGAACGATCCTTCTTTCATCGGTATCTTTTTCAGTAAATGTAGAATCGGTAACCTGAGGGACGGAGATATTGATCAAAGAGACCCTGGAGTGTACTTTAGGATGGAA
AGGGATCTCAGAAATAGGCAGGACAGGATGGGAGTGGAGATCAGAGAACGACTGAAATGGGATATAAGGGAAGGGGTGATAGGGTCAGAAACATTCAATCCCAAACA
CGCCATTATCGTCACATGGAAAAATATCTCTTTCAATGGAGGTTTTGGCAATGCTCTCTACCAGACTAACACTTTCCAAATGATCCTCGCCACTGATGAAGTTTTCA
CCTACGCCATGTTCAACTACTTGAATCTTGACTGGACCACCCACACTGAAGCGGGAGGCGACACAAGAAAAGGAGAAGGAGGAGTTCCCGCTTTTGTGGGATTCAAC
GCTGGAAACGGTACTAGAAGTTTTGAATACAAACCATACAGTCAAGAATCTGTTATTCGAGATCTCACACAAACTGGTTTCGCTAATGGTTTCAAAGGAAGGCACAT
TTTCCGAATCGACGAAAATATCCTAACTGGAACATGCAATAAAGATATAGATGGTGCTAATCTACCGTTAATGATATCTCCAGAAAGTGGAAATATGCTGGGTGGAA
CAATAGTGAATATAACAGGACCTTGTTTCGGCCTAGACGACCAAGTTAAATGCAAATTTGATGTAGCCGATGAAATAAATGGCGTCGTTATAGATAAAAACAGGGCT
ATATGCATCCAACCTAGACTGTATGCCGAAGGATGGGTGAATTTACAAATAGCCATAGGGGCTGGGGTATACAAATGGAAGGGAAAATATTATGTCGAATCCCCCGC
AGCGGCATCTCAAAAAATCTACTTCAAGGACATGAAGGTTCATGAAAAATCGCCTAGTGAAATAAGAATAACTTGGGAAAAATACAACTTGACCACTAACGAAAACG
CTAACATTCGCATCTCCTTATGGGGTTACAGAGAAACAACAATAAGGCCAACGTTCGTTTACATCACTGATATCGCAGACAGTCTCCAAAATACTGGAGAGTATACC
ATCGTACCGTCCCAATATAGAACAAAAGTTAATGAGTTTCTCACGGATATCAAATTTGGTTTCTTGCAAATTAACTTGACTGAATCTATCAAAGTGAACACTTATAC
ATCTGTACAACGATCAGTGGAAATAGTTCCTGTTGTATGGAGTCGACCCATTCCCCTAGGATGGTACTTTCAGTTCCAATGGGAAAATATGTATGGACGAAGCTGGC
CCAAAGCACTCTGCGATGACTGGCTAAGAACAGACAGATACCTGAAAAACTTTGCTCATGAGTTACCTCAATGCCCTTGCACTGTAGAACAGGCTTTGGCAGACAAA
GGGAGGTATATGCCCGACTTTGATTGCGACAAGGACTCAAATCCCGTATGCTACTACAATAACCAAGCTCTGCACTGTGTGAAAACAGGATCACCAACGTTGGAGGG
ATCAGAACAGCAGTGCTGCTATGACAAAAACGGGTATCTCATGTTATCATACGATCAGCAGTGGGGTTCAAGTCCACGGCGTTGCCACAATCTGGGAAAAATGCCCT
ACAACGAAGCAACAAAAGTTCCAACCTTATCGCAATGGTTCAACGATATGGTACCGAAGTATCTTTGCTGTTTGTGGCAGGAAGAACAGGCGGTGGGTTGCGAAACG
CTGAGATTCGAAAGAAGACCAACTCAGGACTGTGTCGCGTACCAAGCTCCAGGGATTGCTGGGATTTACGGAGATCCCCACGTCATCACTTTCGATGACGTCGAGTA
CACCTTCAACGGGAAAGGAGAGTTTGCTCTTGTGAAATCTGTGACACAAACTGACAACTTGGAGGTGCAAGGCAGATTTGAGCAAATGGACCCTAACGCCTACGGAG
AAGTACGTGCAACACAACTGACTTCAATTGTGGCAAGGGGAAACAACACCATAGCAGTGGAGGTCAGAAGGAGGCCCTTGGATGCTAGGTGGAGGTATAGGCTGGAT
GTCATTGCTGATAATAGGAAGTTGTTCTTCGACAGACCCTCTTTGAAATTCCAACATTTCCAAGGAGTGACTATTTATACACCTACTTATATCCTCAATCAGTCTGA
AGTCATCATTATGTTTGATAACGGAGCAGGAGTTCAAGTAATGGATAACCAGGGATTCATGACCGCGAGGGTGTATCTTCCTTGGTCATTCATCAACAAAACTGTTG
GTCTCTTTGGCAACTGGAGTTTCAATAAGGAAGATGACTTCACTCTTCCTGATGAGTCGAAGGCTGCCGTCGTGGGTAATATCAATGATATGGAAAGGGTCTACAAC
GATTTTGGTTCCAAATGGATGGTGGACGACGTACTAGATCCGAAAAGAGGTAGATCCCTATTTTTCAGAGAATTCGGCAGATCATCGGCAACGTACAACAACAAAAC
TTTCAAACCGCAGTTCCTTATGTTACCTGAGGACATAATACCCGCAAACAGGTCGATACAGATACAGAGAACTTACGACATTTGTAGCACAAAAATGTACGAATGCT
ACTACGATTATGCCATGACGCTCAACAGAGATCTTGCCCATTTACTCAGAATTATAAAGCAACCATATATCAACTCAAAGAAACGACGAGGCAGAAGGTTGTTTCT
TGCGGAGTTCTGGAAACACCGCGATTCGGTAGGAAGAGTACTTTTCTTTTTATACCAGGAACCAAAGTCACTTACGAGTGCAATCAAGACTTCGTATTGGTGGGAGA
TCCCAGAAGAGAATGTTTGGCAGATGGCACATGGAATGCTCCTGAATATGGCTACACCGAATGTTTACGTCAACAAGAATATTCTTCCCGAACCGCCATGATCACTT
GGTCCATAATCCTTGCAGTTCTTATACCATTAATCCTATTGATACTTTTCGCGGGATACAAAGTGTATCAAAAAATCAAAGGCGACTCATGGGAGGACAACGATTCA
ACTAAACCTAAAAAATTGCAAGCCTTCAATCGGGCACTCAGTCCATACCAAGATGAAGAAGACGACGATGATGACGACTATGTTCCCAATCCCAGTGACAAGTCAGA
AAGTCAACAAGAATATTCTCAGCGCCAATCAGCCATTGCCTCTGGAGCCGTTCTCGCAATAATTATTCCACTAGTTTTATTATTTGTATATCTGGCTTATATGTTCC
TCAAGAAGAAACAGAAAGAACGAGACGAAGAAAATTTACAAACGCAAGCGTACGAGCAACAGAAAAGACAAGCTCAGGAAGCTGCTGCTAGAAAATTAACTGCTGCC
GAAGAGTACAACTCAGATGAAGACGATAATAACAGCAACGTTACGAGCACAGCAAAAGAAACAACAGTGTATTAGTATTAGATATGGTAACACTTTTACGATATTAT
TATTTTGGTTGGCATACTTCTCAGACAAACAGTCCAAATTATTATTTGCAACAAGAACTATTTGAATTTCGAAGTTTCCTTTGAAGATCCCAAGTTGCTTATTATCC
TAAATTTTTGCAAAATGTCAAAATCGTACTATTCCATCAGTTGTATATTTTATTTTTCTGAGATAGTCAGGCCAACAGGTATCTAAAGGAGTGAATAACAGATTTTT
CATATTTTCCAGTTTTATCTGATTCAATAATGCACTATGGGTTTTTTCTGTTAATGGTTTTTATAACACCTCTGTTGCAAAATACATATTAGGTAATATAAATATAC
GTAATTAAGTCGTAATTAAGTATTATAACAAAATGGCGTTCTCTATTATTTAAATTGTAATTAGATATATATTTTTTATATTCATAATCACTTTTGCATGTAAAGTA
GGAGTCCAATTTTCTGTACTTCAACCATCACTATCGCTACTTACCCCGATCACTTTTTGGATGCACATTTCATGCACATTTTTGCACATTGTTTGCACAGTAAGTTC
ACTTTGCAAACAATGTGCGAAGACTATGCGGGAGGACATGTTTATGCAGTAATTTTGTATATATGTGAGTAGGTATATACCTACTATTTTCATCATACTCTGGTCCT
ACTTTTCTTATTTACAGTGATAAGATGATGTTTGAAATCTTGAATTAGGTACAATGAATTCATTTTTGTCAGTTATTTGAAACATTTTCGGATATTTTAATAAGACT
GTGTTTTATCAAAGTTCAGTTTGATTGTCCAAATACTAGTAGATCATTAGAGTCCTACTGTAGTTTTAGCATTTTAATAAAGTAATTCAATATTAATATACAAGTAA
TCTTGCACTTCTGCTAACTCAAAATTTTGCAAAATTATTCACTCGAATCAGTATTCACACGTTTCTCTTTCAGAAATAACCAAAATATTTCACCCCATACACAAATT
TCACTAAACCTAATGAAAATATATATTCTCCAATAAATTTTTTGAAAACAAGATG

Sequence features legend:

Exon11
dsMESH fragment cloning primers
pRCR primers (listed in Supplementary Table 1)
qPCR probe (listed in Supplementary Table 1)

##### 1.2 dsRNA sequences used in the trials

>dsMESH (417 bp)
CTTTTGTGGGATTCAACGCTGGAAACGGTACTAGAAGTTTTGAATACAAACCATACAGTCAAGAATCTGTTATTC
GAGATCTCACACAAACTGGTTTCGCTAATGGTTTCAAAGGAAGGCACATTTTCCGAATCGACGAAAATATCCTAA
CTGGAACATGCAATAAAGATATAGATGGTGCTAATCTACCGTTAATGATATCTCCAGAAAGTGGAAATATGCTGG
GTGGAACAATAGTGAATATAACAGGACCTTGTTTCGGCCTAGACGACCAAGTTAAATGCAAATTTGATGTAGCCG

```
ATGAAATAAATGGCGTCGTTATAGATAAAAACAGGGCTATATGCATCCAACCTAGACTGTATGCCGAAGGATGGG
TGAATTTACAAATAGCCATAGGGGCTGGGGTATACAAATGGA

>dsEGFP (423 bp)
CCACAAGTTCAGCGTGTCCGGCGAGGGCGAGGGCGATGCCACCTACGGCAAGCTGACCCTGAAGTTCATCTGCAC
CACCGGCAAGCTGCCCGTGCCCTGGCCCACCCTCGTGACCACCCTGACCTACGGCGTGCAGTGCTTCAGCCGCTA
CCCCGACCACATGAAGCAGCACGACTTCTTCAAGTCCGCCATGCCCGAAGGCTACGTCCAGGAGCGCACCATCTT
CTTCAAGGACGACGGCAACTACAAGACCCGCGCCGAGGTGAAGTTCGAGGGCGACACCCTGGTGAACCGCATCGA
GCTGAAGGGCATCGACTTCAAGGAGGACGGCAACATCCTGGGGCACAAGCTGGAGTACAACTACAACAGCCACAA
CGTCTATATCATGGCCGACAAGCAGAAGAACGGCATCAAGGTGAACTT

>dsGFP (966 bp)
GACTCCTATAGGGAGACCGGCAGATCTGATATCACAAGTTTGTACAAAAAAGCAGGCTCCATGAGTAAAGGAGAA
GAACTTTTCACTGGAGTTGTCCCAATTCTTGTTGAATTAGATGGTGATGTTAATGGGCACAAATTTTCTGTCAGT
GGAGAGGGTGAAGGTGATGCAACATACGGAAAACTTACCCTTAAATTTATTTGCACTACTGGAAAACTACCTGTT
CCATGGGTAAGTTTAAACATATATATACTAACTAACCCTGATTATTTAAATTTTCAGCCAACACTTGTCACTACT
TTCTGTTATGGTGTTCAATGCTTCTCGAGATACCCAGATCATATGAAACGGCATGACTTTTTCAAGAGTGCCATG
CCCGAAGGTTATGTACAGGAAAGAACTATATTTTTCAAAGATGACGGGAACTACAAGACACGTAAGTTTAAACAG
TTCGGTACTAACTAACCATACATATTTAAATTTTCAGGTGCTGAAGTCAAGTTTGAAGGTGATACCCTTGTTAAT
AGAATCGAGTTAAAAGGTATTGATTTTAAAGAAGATGGAAACATTCTTGGACACAAATTGGAATACAACTATAAC
TCACACAATGTATACATCATGGCAGACAAACAAAAGAATGGAATCAAAGTTGTAAGTTTAAACATGATTTTACTA
ACTAACTAATCTGATTTAAATTTTCAGAACTTCAAAATTAGACACAACATTGAAGATGGAAGCATTCAACTAGCA
GACCATTATCAACAAAATACTCCAATTGGCGATGGCCCTGTCCTTTTACCAGACAACCATTACCTGTCCACACAA
TCTGCCCTTTCGAAAGATCCCAACGAAAAGAGAGACCACATGGTCCTTCTTGAGTTTGTAACAGCTGCTGGGAAT
ACACATGGCATGGATGAGACCCAGCTTTCTTGTACAAAGTGGTGAATATCAGCTTATCGATACCGT
```

##### 1.3 Evaluation of *in vivo* dsRNA production

We estimated the *in vivo* production of dsMESH to 10 µg per ml of culture. Isolated RNA was treated with DNase I to remove residual DNA and with RNase $I_f$ that preferentially degrades single stranded RNA. Whereas most other bands RNA were degraded by RNase, bands corresponding to dsGFP and dsMESH remained visible (Supplementary Figure 2).

##### 1.4 Feeding trial one: continuous treatment, 2nd instar larvae

This trial was performed June-July 2016 using *in vitro* synthesised dsMESH and dsEGFP. Water treatment was used as blank control and dsEGFP as a non-specific dsRNA (negative control). Forty beetles were selected randomly for each treatment and were reared from 2nd larval instar till pupation on treated detached leaves that were exchanged daily. Larval weight, survival, pupation duration, adult emergence, and gene silencing efficiency were measured daily until (Supplementary Figure 4). Gene silencing determined by qPCR showed 71% reduction of *mesh* expression after 4 days of treatment (whole larvae were sampled). Treatment with dsMESH resulted in 87.5% mortality by 5th day of treatment and 100% larval mortality by 8th day (Supplementary Figure 4).

In this trial we observed an unusually high mortality rate (approx. 40%) for dsEGFP and water control treatments (Supplementary Figure 4). The reason for this might have been CPB egg stress induced by transport or environmental differences between laboratories.

##### 1.5 Feeding trial two: continuous treatment, 4th instar larvae

In trial two, performed June-July 2016, three potted potatoes per treatment were sprayed, placed in a glass container in the laboratory greenhouse, and infested with early 4th instar CPB larvae. These larvae were previously reared on non-treated potato foliage. Larval mortality was checked 5 dpt (one dead larva in water and dsMESH treatment groups, two dead larvae in dsEGFP group) and 12 dpt (only one additional dead larva in dsMESH treatment group).

Adult emergence from the plant substrate was observed until 22 dpt, then the substrate was inspected for beetle carcasses. The phenotypes of emerged adults and carcasses recovered from the substrate were photographed (Supplementary Figure 5).

##### 1.6 Feeding trial three: discontinuous leaf disk treatment with dsRNA, 2nd instar larvae

Trial three was performed in December 2016. Individual potato leaf disks, measuring 10 mm in diameter, were placed in wells of a 24-well plate with preloaded 0.5% agarose gel. *In vitro* synthesised dsRNA (conc. 0.5 µg/µl) was pipetted onto the leaf disks (3x 0.5 µl for the first treatment). The droplets were left to dry at room temperature for approx. 1 hour. Individual 2nd instar larvae were first starved for 2 hours, then placed into the wells to feed on the leaf disks overnight. After treatment, the larvae were transferred to non-treated potato foliage and their weight and mortality was recorded daily until 7 dpt.

##### 1.6 Feeding trial four: discontinuous leaf disk treatment with in vitro and in vivo synthesised dsRNA, 2nd instar larvae

Trial four was performed in April 2018. Treatments were performed similarly as in trial three, but instead of using *in vitro* synthesised dsRNAs, supernatants of dsRNA producing bacterial lysates (conc. 100 ng/µl) were pipetted onto leaf disks measuring 7.5 mm in diameter. Treatments with 60-fold, 600-fold, and 6000-fold serial dilutions of *in vitro* synthesised dsMESH (6, 60, and 600 ng per leaf disk, respectively) were used as comparison to evaluate the potency of *in vivo* synthesised dsMESH. After treatment, the larvae were transferred to non-treated potato foliage and their weight and mortality was recorded daily until 10 dpt.

##### Feeding trial five: egg treatment

Sixty CPB egg masses laid within one day were collected from the laboratory colony and randomly assigned to the three treatment groups. Egg spraying (Supplementary Figure 6) was performed with *in vitro* synthesised dsRNAs (conc. 0.5 µg/µl). Most larvae hatched three days after treatment. The 1st instar larvae were transferred to untreated detached potato leaves that were exchanged daily. Larval mortality was recorded daily until 13 dpt.

##### Survival analysis statistics for laboratory feeding trials

The following are text format statistical outputs of the R survival package. For each trial, the R function call with parameters is given and significant results are marked with red text colour.

*Feeding trial one: continuous treatment, 2nd instar larve (June-July 2016)*

Whole trial

```
Call:
coxph(formula = Surv(june2016.data[, "time"], june2016.data[, "event"], type = "right") ~ june2016.data[, "treatment"])
  n= 240, number of events= 178
                                                        coef exp(coef) se(coef)     z Pr(>|z|)
june2016.data[, "treatment"]2_dsEGFP (neg. control)   0.2041    1.2265   0.2865 0.712   0.4762
june2016.data[, "treatment"]6_dsMESH                  2.7909   16.2964   0.3145 8.874  < 2e-16 ***
---
Signif. codes:  0 '***' 0.001 '**' 0.01 '*' 0.05 '.' 0.1 ' ' 1

                                                     exp(coef) exp(-coef) lower .95 upper .95
june2016.data[, "treatment"]2_dsEGFP (neg. control)      1.226    0.81535    0.6995     2.151
june2016.data[, "treatment"]6_dsMESH                    16.296    0.06136    8.7978    30.187

Concordance= 0.714  (se = 0.026 )
Rsquare= 0.376   (max possible= 0.999 )
Likelihood ratio test= 113.3  on 5 df,   p=0
Wald test            = 109.8  on 5 df,   p=0
Score (logrank) test = 146.7  on 5 df,   p=0
```

Larval stage only (until 14th day of trial)

```
Call:
coxph(formula = Surv(june2016.data[, "time"], june2016.data[, "event"], type = "right") ~ june2016.data[, "treatment"])
  n= 240, number of events= 143
                                                         coef exp(coef) se(coef)     z Pr(>|z|)
june2016.data[, "treatment"]2_dsEGFP (neg. control)   0.02204   1.02229  0.35440 0.062   0.9504
june2016.data[, "treatment"]6_dsMESH                  2.65720  14.25635  0.33037 8.043 8.88e-16 ***
---
Signif. codes:  0 '***' 0.001 '**' 0.01 '*' 0.05 '.' 0.1 ' ' 1

                                                     exp(coef) exp(-coef) lower .95 upper .95
june2016.data[, "treatment"]2_dsEGFP (neg. control)      1.022    0.97820    0.5104     2.048
june2016.data[, "treatment"]6_dsMESH                    14.256    0.07014    7.4609    27.241

Concordance= 0.723  (se = 0.028 )
```

```
Rsquare= 0.368   (max possible= 0.997 )
Likelihood ratio test= 110.3  on 5 df,   p=0
Wald test            = 107.3  on 5 df,   p=0
Score (logrank) test = 144.2  on 5 df,   p=0
```

*Feeding trial two: continuous treatment, 4th instar larvae adult emergence data (June-July 2016)*

```
Call:
coxph(formula = Surv(adult_emerg.data[, "time"], adult_emerg.data[, "event"], type = "right") ~ 
adult_emerg.data[, "treatment"])

  n= 85, number of events= 47 

                                           coef exp(coef) se(coef)      z Pr(>|z|)    
adult_emerg.data[, "treatment"]2_dsEGFP  0.7208    2.0560   0.3060  2.355 0.018505 *  
adult_emerg.data[, "treatment"]3_dsMESH -2.3126    0.0990   0.6177 -3.744 0.000181 ***
---
Signif. codes:  0 ‘***’ 0.001 ‘**’ 0.01 ‘*’ 0.05 ‘.’ 0.1 ‘ ’ 1

                                        exp(coef) exp(-coef) lower .95 upper .95
adult_emerg.data[, "treatment"]2_dsEGFP     2.056     0.4864   1.12862    3.7454
adult_emerg.data[, "treatment"]3_dsMESH     0.099    10.1006   0.02951    0.3322

Concordance= 0.801  (se = 0.033 )
Rsquare= 0.402   (max possible= 0.989 )
Likelihood ratio test= 43.76  on 2 df,   p=3e-10
Wald test            = 25.27  on 2 df,   p=3e-06
Score (logrank) test = 41.1  on 2 df,   p=1e-09
```

*Feeding trial three: treatment with dsRNA twice (December 2016)*

```
Call:
coxph(formula = Surv(dec2016.data[, "time"], dec2016.data[, "event"], type = "right") ~ dec2016.data[, "treatment"])
  n= 80, number of events= 25 
                                       coef exp(coef) se(coef)     z Pr(>|z|)    
dec2016.data[, "treatment"]2_dsEGFP  0.7298    2.0746   1.2248 0.596 0.551284    
dec2016.data[, "treatment"]5_dsMESH  3.9675   52.8518   1.0479 3.786 0.000153 ***
---
Signif. codes:  0 ‘***’ 0.001 ‘**’ 0.01 ‘*’ 0.05 ‘.’ 0.1 ‘ ’ 1

                                    exp(coef) exp(-coef) lower .95 upper .95
dec2016.data[, "treatment"]2_dsEGFP     2.075    0.48203    0.1881     22.88
dec2016.data[, "treatment"]5_dsMESH    52.852    0.01892    6.7777    412.13

Concordance= 0.832  (se = 0.063 )
Rsquare= 0.434   (max possible= 0.928 )
Likelihood ratio test= 45.53  on 4 df,   p=3e-09
Wald test            = 44.29  on 4 df,   p=6e-09
Score (logrank) test = 79.01  on 4 df,   p=3e-16
```

*Feeding trial four: testing in vivo synthesised dsRNAs (April 2018)*

**all – coxph**

```
> fitcox <- coxph(Surv(april2018.data[, "time"], april2018.data[, "event"], type='right') ~ april2018.data[, "treatment"])
Warning message:
In fitter(X, Y, strats, offset, init, control, weights = weights,  :
  Loglik converged before variable  1,2,3,4,5 ; beta may be infinite. 
> summary(fitcox)
Call:
coxph(formula = Surv(april2018.data[, "time"], april2018.data[, "event"], type = "right") ~ april2018.data[, "treatment"])

  n= 143, number of events= 97 
```

```
                                                         coef exp(coef)  se(coef)     z Pr(>|z|)
april2018.data[, "treatment"]2_dsGFP_E_coli          1.782e+01 5.483e+07 3.485e+03 0.005    0.996
april2018.data[, "treatment"]3_dsMesh_E_coli         2.110e+01 1.454e+09 3.485e+03 0.006    0.995
april2018.data[, "treatment"]4_dsMesh_in_vitro_6000x 2.022e+01 6.068e+08 3.485e+03 0.006    0.995
april2018.data[, "treatment"]5_dsMesh_in_vitro_600x  2.124e+01 1.675e+09 3.485e+03 0.006    0.995
april2018.data[, "treatment"]6_dsMesh_in_vitro_60x   2.109e+01 1.445e+09 3.485e+03 0.006    0.995

                                                     exp(coef) exp(-coef) lower .95 upper .95
april2018.data[, "treatment"]2_dsGFP_E_coli          5.483e+07  1.824e-08         0       Inf
april2018.data[, "treatment"]3_dsMesh_E_coli         1.454e+09  6.876e-10         0       Inf
april2018.data[, "treatment"]4_dsMesh_in_vitro_6000x 6.068e+08  1.648e-09         0       Inf
april2018.data[, "treatment"]5_dsMesh_in_vitro_600x  1.675e+09  5.972e-10         0       Inf
april2018.data[, "treatment"]6_dsMesh_in_vitro_60x   1.445e+09  6.920e-10         0       Inf

Concordance= 0.822  (se = 0.041 )
Rsquare= 0.65   (max possible= 0.998 )
Likelihood ratio test= 150.1  on 5 df,   p=<2e-16
Wald test            = 44.91  on 5 df,   p=2e-08
Score (logrank) test = 126.4  on 5 df,   p=<2e-16
```

Because the Cox-PH model resulted in degenerate estimate in the group with no events (see the result shaded in grey above), the log-rank statistics of most relevant comparisons was used instead:

**3_dsMesh_E_coli vs 2_dsGFP_E_coli (***)**

```
Call:
coxph(formula = Surv(subst[, "time"], subst[, "event"], type = "right") ~ as.vector(subst[, "treatment"]))
  n= 47, number of events= 28
                                                coef exp(coef) se(coef)     z Pr(>|z|)
as.vector(subst[, "treatment"])3_dsMesh_E_coli  3.4295   30.8608   0.6272 5.468 4.54e-08 ***
---
Signif. codes:  0 ‘***’ 0.001 ‘**’ 0.01 ‘*’ 0.05 ‘.’ 0.1 ‘ ’ 1

                                               exp(coef) exp(-coef) lower .95 upper .95
as.vector(subst[, "treatment"])3_dsMesh_E_coli     30.86     0.0324     9.028     105.5

Concordance= 0.834  (se = 0.063 )
Rsquare= 0.635   (max possible= 0.984 )
Likelihood ratio test= 47.33  on 1 df,   p=6e-12
Wald test            = 29.9  on 1 df,   p=5e-08
Score (logrank) test = 47.93  on 1 df,   p=4e-12


4_dsMesh_in_vitro_6000x vs 2_dsGFP_E_coli (***)
Call:
coxph(formula = Surv(subst[, "time"], subst[, "event"], type = "right") ~ as.vector(subst[, "treatment"]))
  n= 47, number of events= 25
                                                        coef exp(coef) se(coef)     z Pr(>|z|)
as.vector(subst[, "treatment"])4_dsMesh_in_vitro_6000x  2.3923   10.9389   0.5555 4.306 1.66e-05 ***
---
Signif. codes:  0 ‘***’ 0.001 ‘**’ 0.01 ‘*’ 0.05 ‘.’ 0.1 ‘ ’ 1

                                                       exp(coef) exp(-coef) lower .95 upper .95
as.vector(subst[, "treatment"])4_dsMesh_in_vitro_6000x     10.94    0.09142     3.682      32.5

Concordance= 0.776  (se = 0.061 )
Rsquare= 0.434   (max possible= 0.977 )
Likelihood ratio test= 26.78  on 1 df,   p=2e-07
Wald test            = 18.54  on 1 df,   p=2e-05
Score (logrank) test = 27.43  on 1 df,   p=2e-07

5_dsMesh_in_vitro_600x vs 2_dsGFP_E_coli (***)
```

```
Call:
coxph(formula = Surv(subst[, "time"], subst[, "event"], type = "right") ~ as.vector(subst[, "treatment"]))
  n= 47, number of events= 28
                                                        coef exp(coef) se(coef)     z Pr(>|z|)
as.vector(subst[, "treatment"])5_dsMesh_in_vitro_600x  4.356    77.948    1.048 4.155 3.26e-05 ***
---
Signif. codes:  0 ‘***’ 0.001 ‘**’ 0.01 ‘*’ 0.05 ‘.’ 0.1 ‘ ’ 1

                                                      exp(coef) exp(-coef) lower .95 upper .95
as.vector(subst[, "treatment"])5_dsMesh_in_vitro_600x     77.95    0.01283     9.985     608.5

Concordance= 0.839  (se = 0.064 )
Rsquare= 0.667   (max possible= 0.984 )
Likelihood ratio test= 51.64  on 1 df,   p=7e-13
Wald test            = 17.26  on 1 df,   p=3e-05
Score (logrank) test = 49.31  on 1 df,   p=2e-12

6_dsMesh_in_vitro_60x vs 2_dsGFP_E_coli (***)
Call:
coxph(formula = Surv(subst[, "time"], subst[, "event"], type = "right") ~ as.vector(subst[, "treatment"]))
  n= 47, number of events= 28

                                                        coef exp(coef) se(coef)     z Pr(>|z|)
as.vector(subst[, "treatment"])6_dsMesh_in_vitro_60x  3.0559   21.2392   0.5728 5.335 9.53e-08 ***
---
Signif. codes:  0 ‘***’ 0.001 ‘**’ 0.01 ‘*’ 0.05 ‘.’ 0.1 ‘ ’ 1

                                                     exp(coef) exp(-coef) lower .95 upper .95
as.vector(subst[, "treatment"])6_dsMesh_in_vitro_60x     21.24    0.04708     6.912     65.26

Concordance= 0.816  (se = 0.057 )
Rsquare= 0.603   (max possible= 0.984 )
Likelihood ratio test= 43.39  on 1 df,   p=4e-11
Wald test            = 28.47  on 1 df,   p=1e-07
Score (logrank) test = 45.83  on 1 df,   p=1e-11
```

##### *Feeding trial five: treatment of CPB eggs (May 2018)*

```
Call:
coxph(formula = Surv(may2018.data[, "time"], may2018.data[, "event"], type = "right") ~ may2018.data[, "treatment"])
  n= 521, number of events= 197
                                        coef exp(coef) se(coef)      z Pr(>|z|)
may2018.data[, "treatment"]2_dsEGFP -0.4497    0.6378   0.4743 -0.948    0.343
may2018.data[, "treatment"]3_dsMESH  4.0571   57.8074   0.3350 12.112   <2e-16 ***
---
Signif. codes:  0 ‘***’ 0.001 ‘**’ 0.01 ‘*’ 0.05 ‘.’ 0.1 ‘ ’ 1

                                    exp(coef) exp(-coef) lower .95 upper .95
may2018.data[, "treatment"]2_dsEGFP    0.6378     1.5678    0.2517     1.616
may2018.data[, "treatment"]3_dsMESH   57.8074     0.0173   29.9824   111.455

Concordance= 0.889  (se = 0.022 )
Rsquare= 0.643   (max possible= 0.99 )
Likelihood ratio test= 537.2  on 2 df,   p=<2e-16
Wald test            = 268.2  on 2 df,   p=<2e-16
Score (logrank) test = 678.4  on 2 df,   p=<2e-16
```

#### 2 SUPPLEMENTARY FIGURES

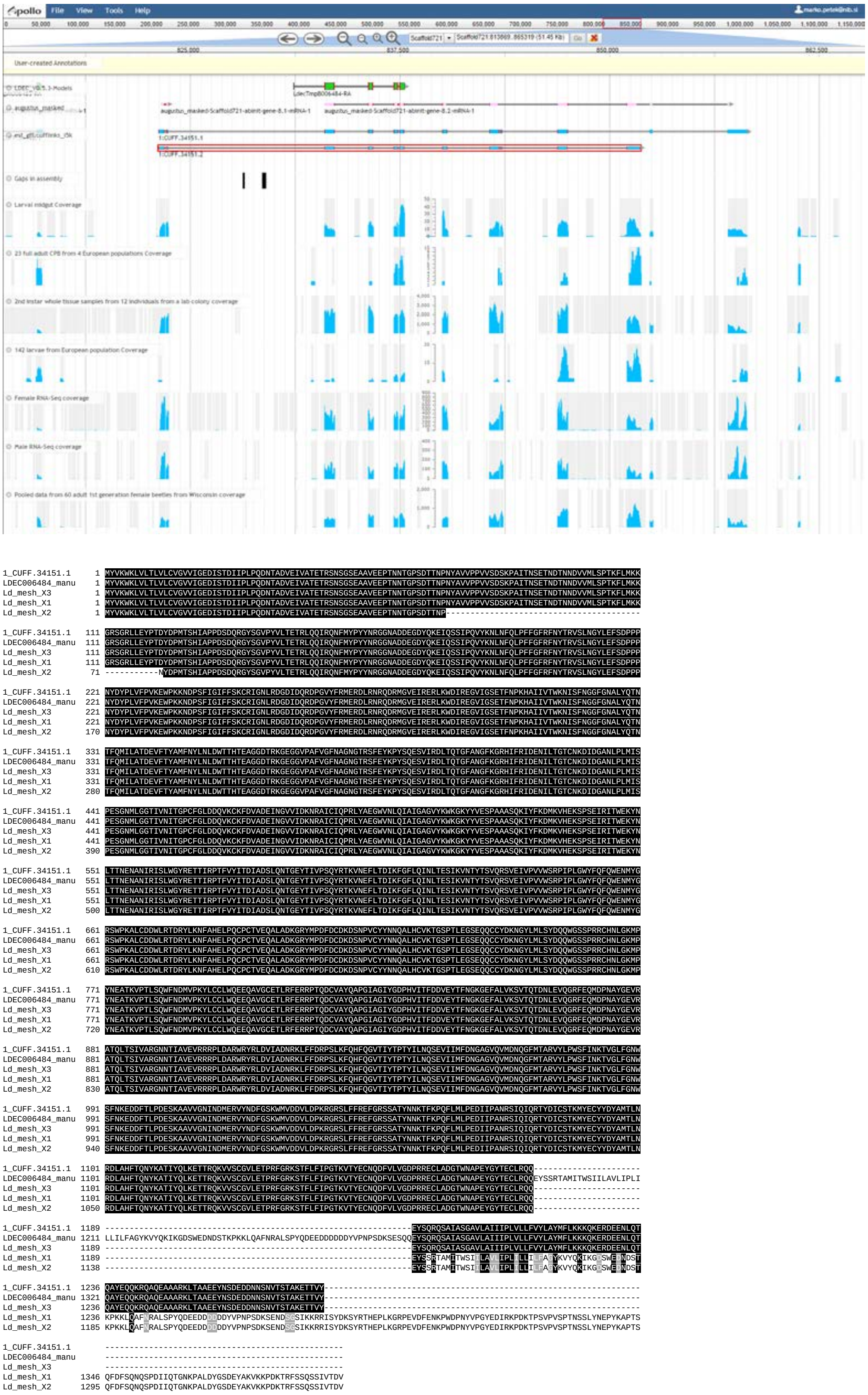

**Supplementary Figure 1. Correction of *mesh* gene model. (A)** WebApollo *mesh* gene models available in i5k CPB genome version 0.5.3. Tracks showing: genome annotation with gene models generated by MAKER, Augustus, and Cufflinks; genome assembly gaps; and RNA-Seq read coverage from different

CPB samples. **(B)** Alignment of CPB Mesh proteins predicted from Cufflinks assembly contig 1:CUFF.34151.1, our manually corrected gene model, and three isoforms (X1-3) predicted by the NCBI genome annotation pipeline (Accessions: XP_023014320.1, XP_023014329.1, and XP_023014336.1, respectively).

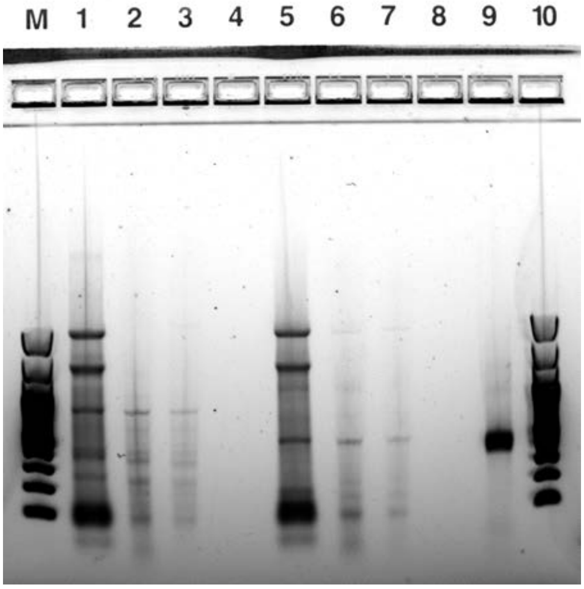


**Supplementary Figure 2. Estimation of *in vivo* synthesised dsRNA amounts.** RNA was run on a 1% agarose electrophoresis gel. Lanes: M - DNA ladder Fermentas 100 bp, 1 - total RNA isolated from *E. coli* producing dsGFP (1 µl loading), 2&3 - RNA from *E. coli* producing dsGFP treated with RNase $I_f$ (loadings of 2 and 4 µl, respectively), 5 - total RNA isolated from *E. coli* producing dsMESH (1 µl loading), 6&7 - RNA from *E. coli* producing dsMESH treated with RNase $I_f$ (loadings of 2 and 4 µl, respectively), 9 - *in vitro* synthesised dsMESH (1 µl loading), 10 - DNA ladder Fermentas 100 bp (1 µl loading).

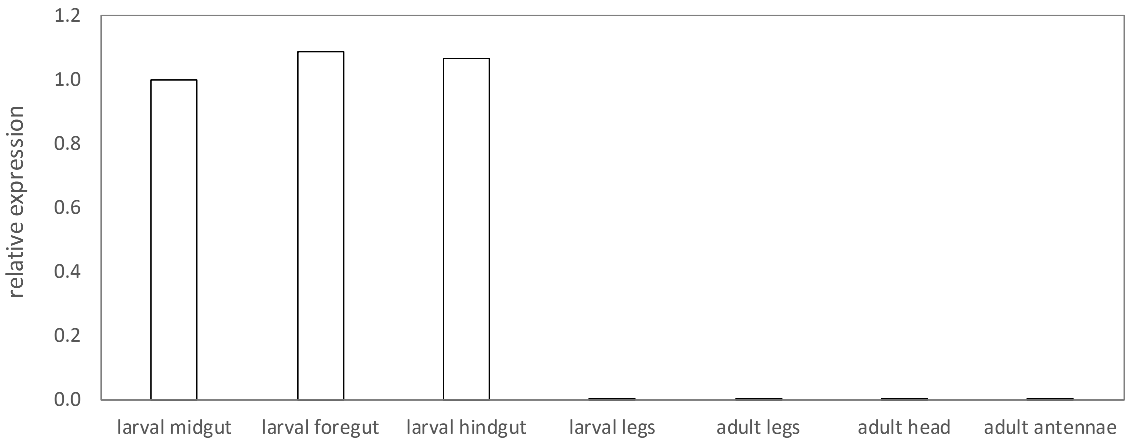


**Supplementary Figure 3. Expression of *mesh* gene in Colorado potato beetle body parts.** Gene expression values are shown relative to midgut. Body parts of 3-4 beetles from same developmental stage were pooled to obtain samples (one biological replicate).

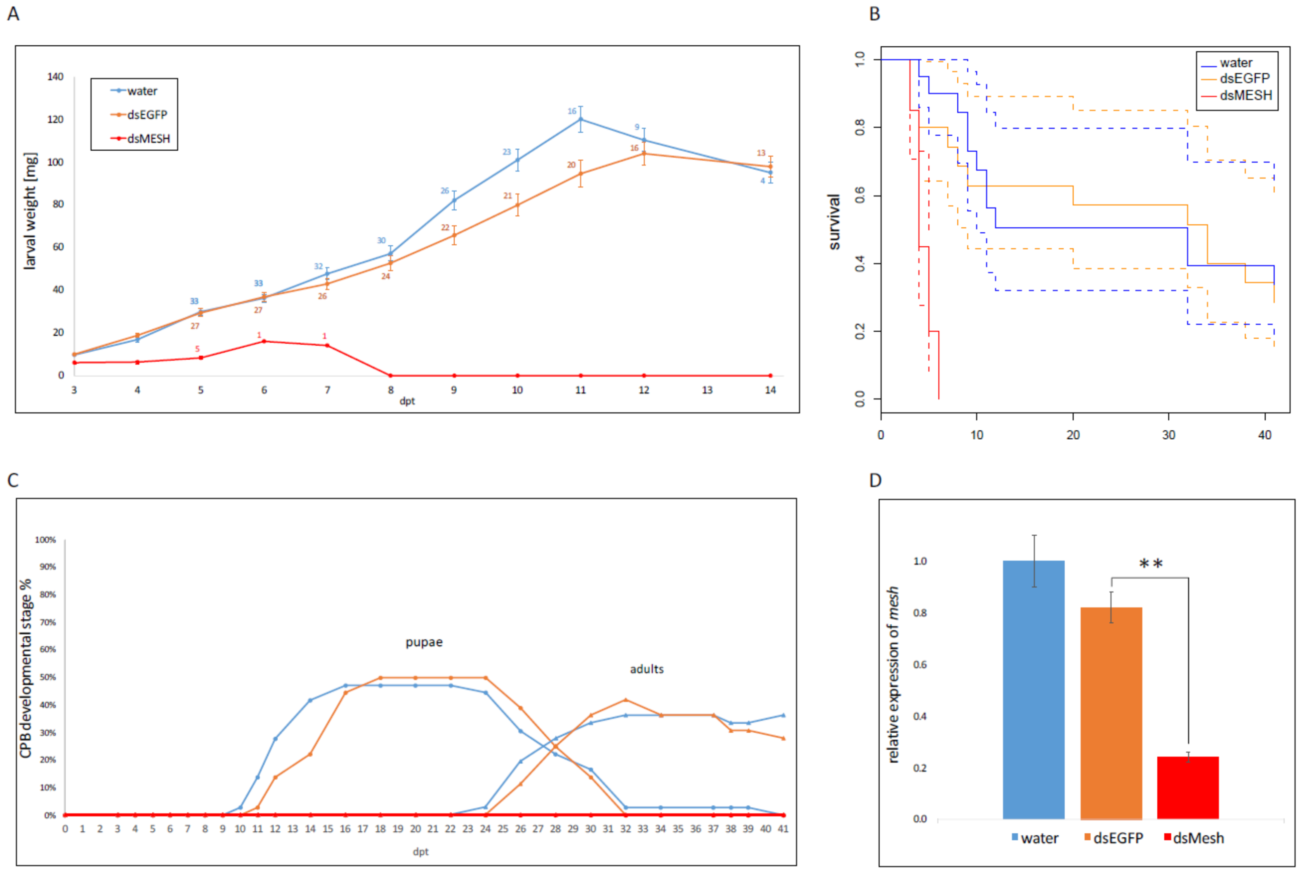


**Supplementary Figure 4. Experimental measurements of continued feeding trial of 2nd instar larvae with *in vitro* synthesised dsRNA (trial one). (A)** Weight of surviving larvae during the treatment. Numbers next to error bars show number of alive larvae for each day. Error bars show standard error of the mean. **(B)** Kaplan-Meier survival curves with 95% confidence interval boundaries (dotted lines). Survival is plotted as proportions. **(C)** Number of pupating (circle) and adult beetles (triangle) throughout the trial. The line colours are the same as for weight measurements. **(D)** Relative target gene expression in whole larvae sampled at 4 days post treatment (dpt). Error bars show standard error of the mean and asterisks denote significant difference in expression compared to dsEGFP treatment ('**' $p < 0.01$).

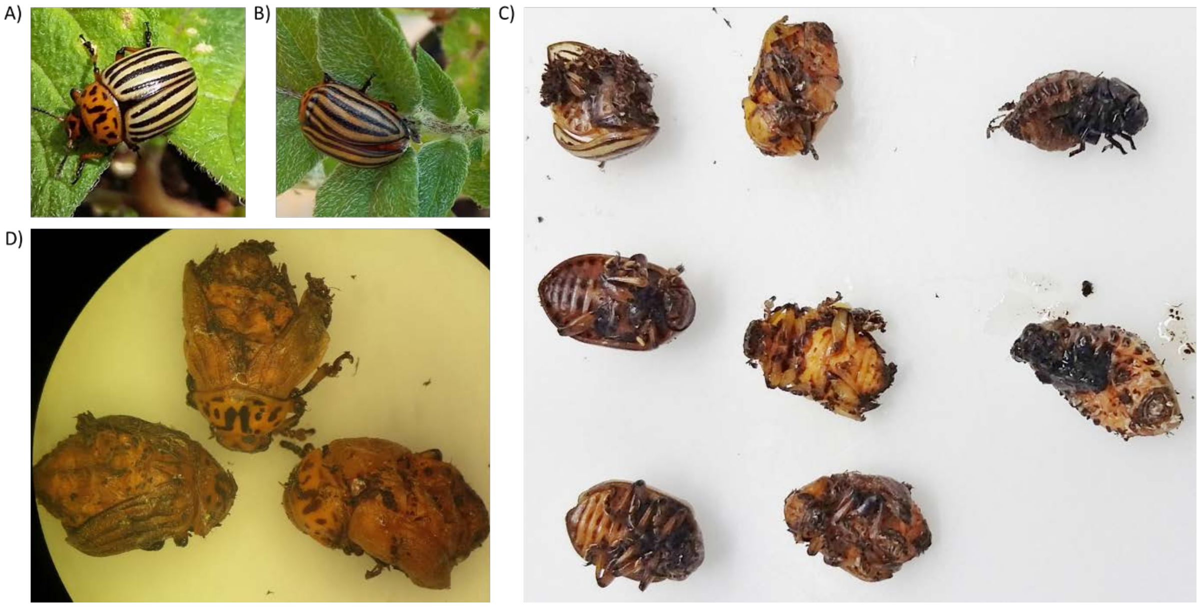


**Supplementary Figure 5. Colorado potato beetle phenotypes observed in adults emerged from 4th instar larvae continuously exposed to dsMESH (trial two). (A)** Normally developed adults emerged from plant substrate from dsEGFP treated beetle group. **(B)** Phenotype of emerged dsMESH treated beetle group. **C)** Six adult and two larval carcasses of dsMESH treated beetle group recovered from plant substrate at the end of trial. D) Backside close-up pictures of three recovered adults.

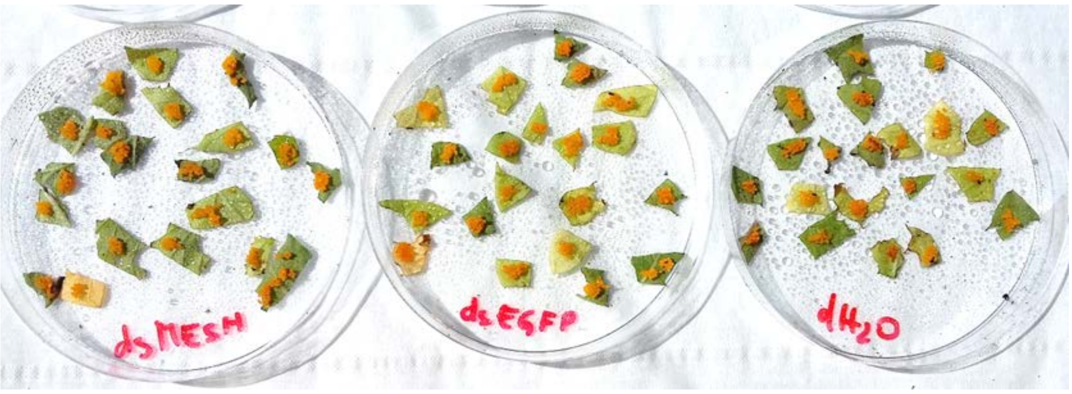


**Supplementary Figure 6. Spraying of Colorado potato beetle eggs (trial five).** Twenty freshly laid egg masses per group were treated and left to dry on air.

#### 3 SUPPLEMENTARY TABLES

**Supplementary Table 1** List of qPCR assays used in this study and their properties according to MIQE guidelines.

[1] NCBI database identifier(s)

[2] FW or F, forward primer; RW or R, reverse primer; P, qPCR probe; NA, not available

[3] Amplification efficiency was calculated from the slope of the log-linear regression curve using the equation $10(-1/\text{slope})^{-1}$

| qPCR assay name | qPCR chemistry | Reference (doi) | Gene ID[1] | Annotation | Primer/probe name[2] | Primer or probe sequence (5'-3') | Primer/probe conc. (nM) | PCR efficiency[3] |
|---|---|---|---|---|---|---|---|---|
| LdMesh | TaqMan | this study | LOC111504068 | mesh, involved in smooth septate junction assembly | LdMesh_qPCR_F | CGGGAGGCGACACAAGAA | 900 | 96% |
| | | | | | LdMesh_qPCR_R | ACCGTTTCCAGCGTTGAATC | 900 | |
| | | | | | LdMesh_qPCR_P | 5'-FAM/AGGAGAAGGAGGAGTTC CCGCTTTTGTG/3'-Zen Iowa BFQ | 250 | |
| 18S | TaqMan MGB | Applied Biosystems | NA | eukaryotic 18S ribosomal RNA (reference gene) | NA | proprietary | proprietary | 92% |
| Ld_smt3 | SybrGreen | Petek et al., 2014 (10.1111/mec.12932) | JN603588 | ubiquitin-like smt3 (reference gene) | con33-F | TACCGATACCCCAACCACATTAG | 900 | 92% |
| | | | | | con33-R | CCAGTTTGCTGTTGGTATACTTCAA | 900 | |
| LdRP4 | SybrGreen | Zhu et al., 2010 (10.1002/ps.2048) | EB761170, KC190033 | ribosomal protein 4 (reference gene) | LdRP4_F | AAAGAAACGAGCATTGCCCTTCCG | 900 | 98% |
| | | | | | LdRP4_R | TTGTCGCTGACACTGTAGGGTTGA | 900 | |
